# Standard Sample Storage Conditions Impact on Inferred Microbiome Composition and Antimicrobial Resistance Patterns

**DOI:** 10.1101/2021.05.24.445395

**Authors:** Casper Sahl Poulsen, Rolf Sommer Kaas, Frank M. Aarestrup, Sünje Johanna Pamp

## Abstract

Storage of biological specimens is crucial in the life and medical sciences. The storage conditions for samples can be different for a number of reasons, and it is unclear which effect this can have on the inferred microbiome composition in metagenomics analyses. Here, we assess the effect of common storage temperatures (deep freezer: -80°C, freezer: -20°C, fridge: 5°C, room temperature: 22°C) and storage times, (immediate sample processing: 0h, next day: 16h, over weekend: 64h, and longer term: 4, 8, 12 months), as well as repeated sample freezing and thawing (2-4 freeze-thaw cycles). We examine two different pig feces and sewage samples, unspiked and spiked with a mock community, and in triplicates, respectively, amounting to a total of 438 samples (777 Gbp; 5.1 billion reads). Storage conditions had a significant and systematic effect on the taxonomic and functional composition of microbiomes. Distinct microbial taxa and antimicrobial resistance classes were in some situations similarly effected across samples, while others were not, suggesting an impact of individual inherent sample characteristics. With an increasing number of freeze-thaw cycles, an increasing abundance of Firmicutes, Actinobacteria, and eukaryotic microorganisms was observed. We include recommendations for sample storage, and strongly suggest including more detailed information in the metadata together with the DNA sequencing data in public repositories to better facilitate meta-analyses and reproducibility of findings.

**IMPORTANCE:** Previous research has reported effects of DNA isolation, library preparation, and sequencing technology on metagenomics-based microbiome composition; however, the effect of biospecimen storage conditions has not been thoroughly assessed. We examined the effect of common sample storage conditions on metagenomics-based microbiome composition and find significant and, in part, systematic effects. Repeated freeze-thaw cycles could be used to improve the detection of microorganisms with more rigid cell walls, including parasites. We provide a dataset that could also be used for benchmarking algorithms to identify and correct for batch effects. Overall, the findings suggest that all samples of a microbiome study should be stored in the same way. Furthermore, there is a need to mandate more detailed information about sample storage and processing published together with the DNA sequencing data at INSDC (ENA/EBI, NCBI, DDBJ) or other repositories.

## Introduction

Metagenomics has emerged as an important technology to reveal microbiome composition at the taxonomic and functional levels, and in the context where they reside such as human and animal hosts and the environment. The sample processing workflow in metagenomics comprises a number of steps, including sample collection, sample storage, DNA isolation, library preparation, DNA sequencing, and data analysis. In microbiome analyses there is an increasing concern about the influence of sample processing on skewing inferred microbial community composition, and which may explain differences observed between studies (1–5). Increasingly, data from different metagenomic projects are combined as part of a study, without taking confounding factors such as differences in sample processing into account. Therefore, it is currently uncertain to what extend it is in fact possible to compare data across studies in a meaningful way without leading to misleading conclusions. Comparisons of data across laboratories and projects are of great importance and could facilitate meta-analyses of human, animal, and environmental microbiome studies, as well as a global metagenomics-based surveillance of pathogens and antimicrobial resistance (6, 7).

A standardization of sample storage conditions can be difficult because sample collection may be performed over a long period of time, suitable storage facilities may be unavailable for instance in field studies, and proper shipping conditions can be limited when samples need to be transported to a central laboratory for analysis (8, 9). A number of studies involving 16S rRNA gene profiling have investigated aspects of storage conditions on the fecal microbiota (10–13). From these studies different variables affecting the microbial composition were identified, such as the temperature and time duration of sample storage, and addition of preservation liquids (10–13). Other factors such as freeze-thawing, atmosphere, UV radiation, and container material might also influence microbial community composition. Some studies have concluded that storage has a limited effect on inferred microbial community composition relative to the variation observed between different samples (10, 11). However, this may depend on how different microbiomes initially were between samples according to certain measures, and the kind of secondary analyses and interpretations that were applied.

The role of storage conditions in metagenomics-based studies is less well described. In a study involving throat swabs, two time points were investigated, namely 0 hours (i.e. DNA isolation immediately upon sample arrival at the lab) and after 24 hours storage at room temperature (5). The study revealed an altering of the microbial community composition and increase in the bacterial ratio to human DNA was observed that was hypothesized to be the result of bacterial growth. More detailed studies are needed to determine the influence of relevant sample storage conditions on inferred microbial community composition of microbiomes.

Here, we assess the effect of storage temperature and time on two different sample types (pig feces and domestic sewage), represented by two samples per same type, respectively. We further investigate the effect of repeated freeze-thaw cycles on inferred microbiome composition. We find that storage has a systematic effect on microbiome composition, resulting from abundance changes of distinct microbial genera. The effect on specific genera was though difficult to generalize across different sample types. A high correlation was observed between patterns of taxonomic microbial community structure and the antimicrobial resistome pattern. Based on our findings we provide recommendations for sample storage conditions.

## Results

In order to determine the effect of relevant microbiome sample storage conditions on different sample types, two pig feces (P1, P2) and two sewage (S1, S2) samples were investigated. These samples were processed and analysed individually as pure sample material, as well as material spiked with a synthetic microbial community composed of eight different microorganisms (Fig. 1A; see also Tables S1, S2 and S3 in the supplemental material). Aliquots of all four samples (P1, P2, S1, S2) were exposed to combinations of 9 different storage conditions: four temperatures (-80°C, -20°C, 5°C, and 22°C) and three storage duration times (0 hours, 16 hours, and 64 hours), and in addition, at the two frozen temperatures (-80°C, -20°C) for three additional storage duration times (4 months, 8 months, 12 months) for samples P1 and S1. Triplicate DNA isolations were performed for each storage situation. This experiment amounted to a total of 304 pig feces and sewage samples. Furthermore, additional aliquots from sample P1 and S1 that were stored at -80°C, -20°C underwent up to four freeze-thaw cycles (Fig. 1A; see also Table S1 in the supplemental material) In addition, 39 control samples were analysed that included DNA isolation blank controls, and the pure mock community. On average, 14.8 million reads were obtained per microbiome sample with a larger output obtained from the pig feces samples (average: 19.9±8.4 million reads) compared with sewage (average: 8.7±3.3 million reads).

**Figure 1:**
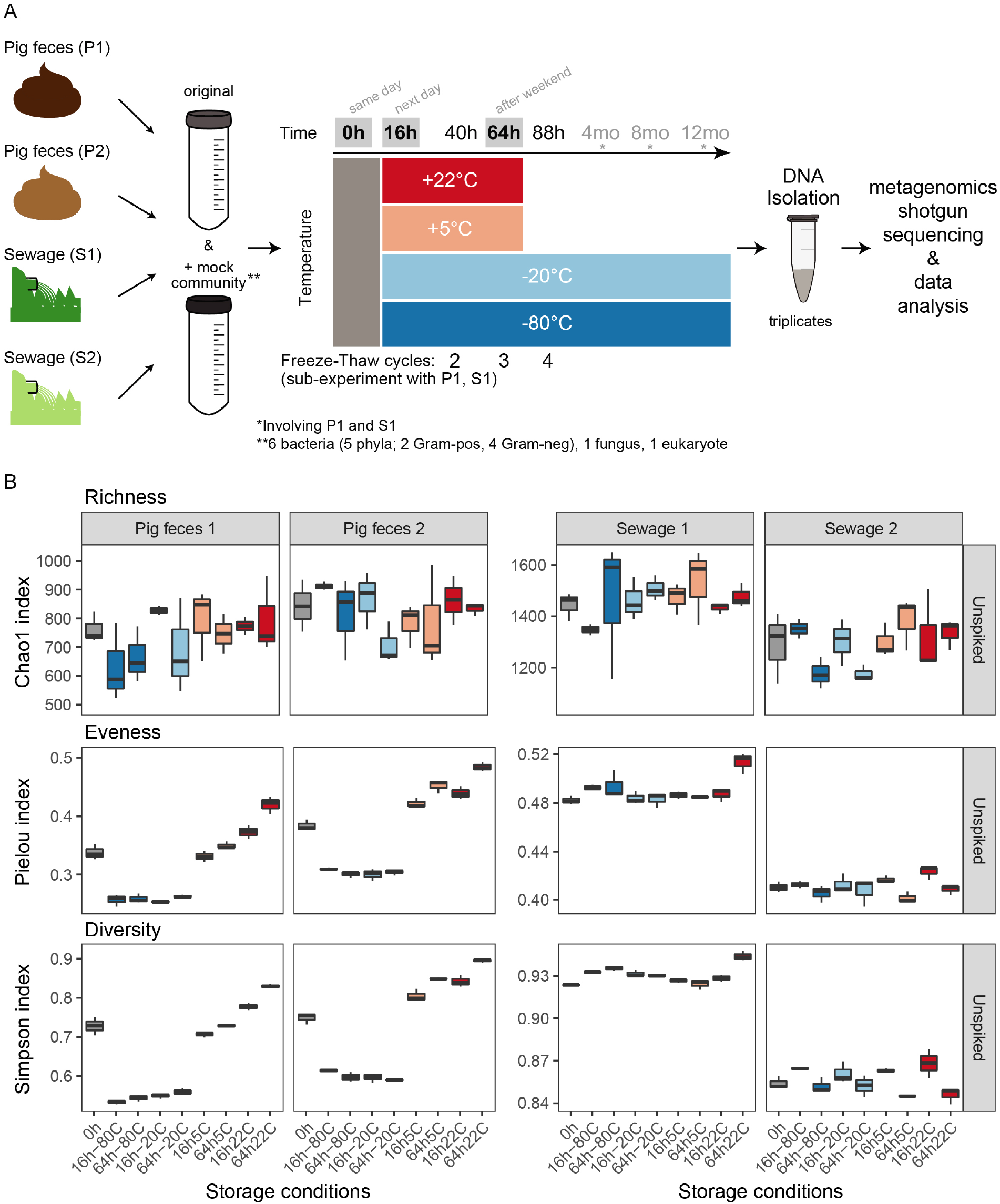
Overview of the experimental setup and alpha diversity of microbiomes. A) Two pig feces and two sewage samples were divided into two large aliquots of which one was spiked with a synthetic mock community composed of eight microorganisms that included two eukaryotes (*Propionibacterium freudenreichii, Bacteroides fragilis, Staphylococcus aureus, Fusobacterium nucleatum, Escherichia coli, Salmonella Typhimurium, Cryptosporidium parvum, Saccharomyces cerevisiae*), respectively. Each of these eight large aliquots was divided into small aliquots which were stored at four different temperatures (-80°C, -20°C, 5°C, and 22°C) for six different durations of time (0 hours, 16 hours, 64 hours, 4 months, 8 months, 12 months). DNA was isolated at the three different time points in triplicates. Additional aliquots for samples P1 and S1 underwent repeated freeze-thaw cycles upon 40 hours, 64 hours, and 88 hours of storage duration. For details, see Materials and Methods. All samples were sequenced on an Illumina HiSeq. B) Alpha diversity was determined: richness (Chao1), evenness (Pielou’s evenness), and diversity (Simpson). The indices were calculated from the count table aggregated at genus level. For the results of the duplicate sets of samples that also harbored the spiked mock community, see Figure S3 in the supplemental material.

Initially, we analysed all samples from all time points (0 hours, 16 hours, 64 hours, 4 months, 8 months, 12 months) together (e.g. see Fig. S1 in the supplemental material). This analysis suggested that inferred microbiome composition differed between storage conditions. However, the reason for some of the patterns were inconclusive, as they may also be related to a technical variation: the samples from the first three time points (0 hours, 16 hours, 64 hours) were processed using another library preparation kit than the samples from the later time points (4 months, 8 months, 12 months) (see Materials and Methods). This prompted us to perform a separate study on assessing the effects of library preparation on inferred microbial community composition involving DNA extracts from the same pig feces and sewage samples from the present study (see (14)). Here, we attempted different strategies of correcting for the underlying batch effects and did not find a way that lead to results we could rely on with confidence. Therefore, we focus our analysis in this study on the first time points (0 hours, 16 hours, 64 hours) as well as the sub-experiment involving a series of freeze-thaw cycles, all undergoing the same library preparation and sequencing strategy (i.e. in all 287 samples). In addition, we make accessible under accession number PRJEB31650 the complete metagenomic datasets of 343 samples presented in this study, as well as 95 samples processed with different combinations of library preparation and sequencing platform with detailed metadata (see Table S1 in the supplemental material). This dataset of 438 samples (total of 777 Gbp, 5.1^10*9 reads) could for example be used to benchmark different algorithms adjusting for batch effects.

### Storage conditions impact on microbial community richness, evenness, and diversity

To assess the level of sequencing depth in relation to the genera observed, rarefaction analyses were performed. The rarefaction curves of individual samples (P1, P2, S1, S2) had similar profiles in which a plateau was approached for most samples, suggesting that the sequencing depth was adequate (see Fig. S2 in the supplemental material).

Overall, the pig feces (P1, P2) and sewage samples (S1, S2) exhibited similar richness (Chao1), eveness (Pielou), and diversity (Simpson) whether they were spiked or not spiked with the mock community, respectively (Fig. 1B, and Fig. S3 in the supplemental material). Richness (Chao1) was similar, independent of storage condition, for the two pig feces samples and two sewage samples, respectively. In contrast, for the pig feces samples the evenness (Pielou) and diversity (Simpson) was lower under frozen storage conditions (-20°C, -80°C), while eveness was higher when samples were stored at positive degree Celsius (+5°C, +22°C), compared to the samples that were not stored and processed immediately (0h) (Fig. 1B, and Fig. S3 in the supplemental material). This was not observed for the sewage samples, for which eveness (Pielou) was similar across all conditions, and slightly increased when stored at 22°C for 64h. This was also reflected in the Simpson’s diversity index that takes both, richness and evenness into account. In summary, the storage conditions had an effect on microbial community evenness (Pielou) and diversity (Simpson) in the pig feces samples, and a lower effect on the sewage samples.

### Samples cluster according to their origin

A pair-wise comparison of all samples was performed by calculating Bray-Curtis dissimilarities on Hellinger transformed data and then visualized using PCoA and Box Plots (Fig. 2A, and see Fig. S4 in the supplemental material). The 212 samples clustered largely according to the sample matrix they originated from (i.e. P1, P2, S1, S2. The two pig feces samples appeared more similar to each other as compared to the two sewage samples, respectively. A separate two-dimensional representation of the pig feces samples (P1 and P2) revealed however that they could be discriminated (Fig. 2B, and see Fig. S5 in the supplemental material). A clear separation of the two sewage samples (S1 and S2) was also observed (Fig. 2C, and see Fig. S5 in the supplemental material). This indicates that the examined storage conditions did not influence the ability to discriminate between the two different pig feces and two different sewage samples, respectively (Fig. 2B and C).

**Figure 2:**
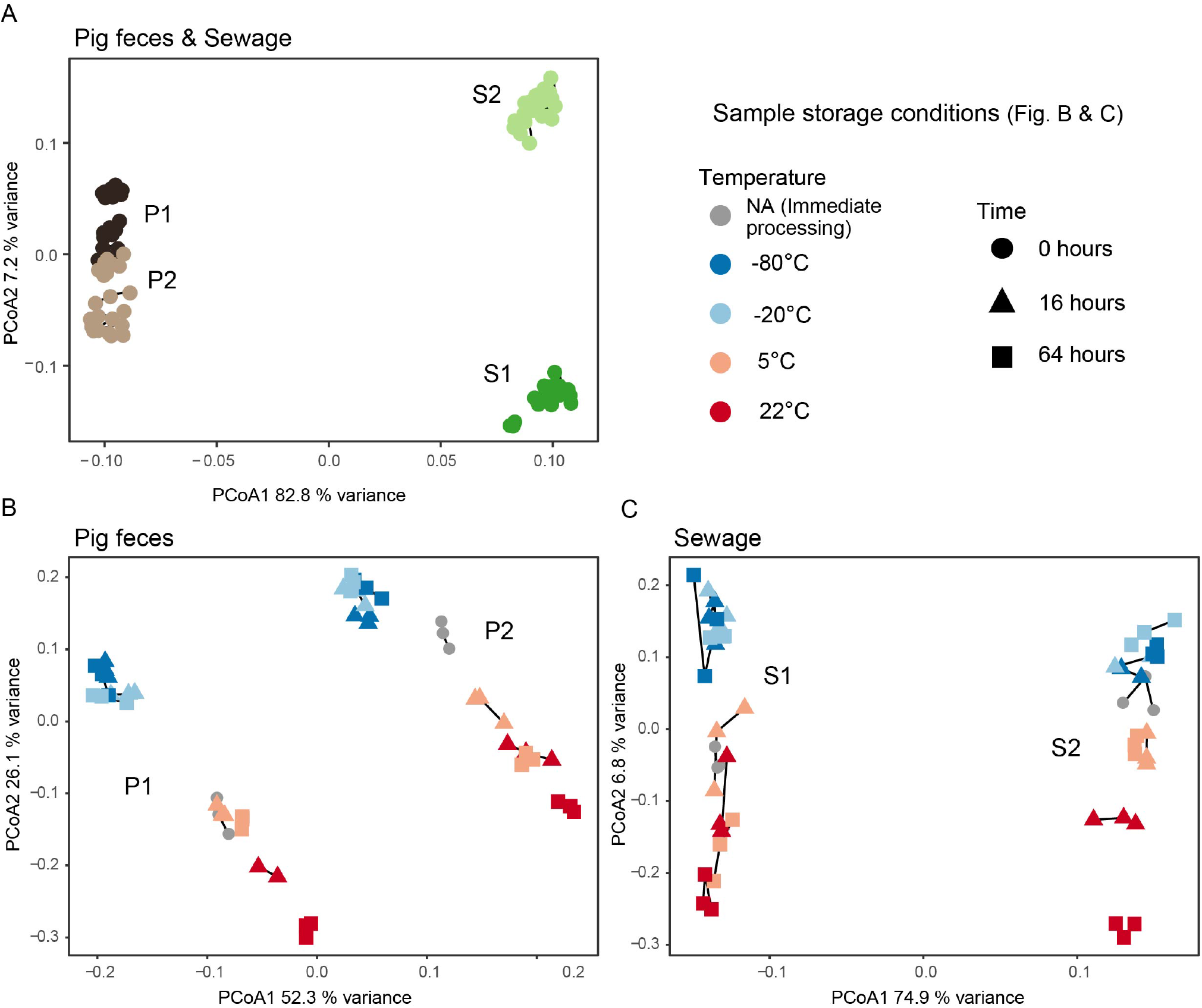
Principal coordinates analysis (PCoA) of all unspiked samples (A), all unspiked pig fecal samples (B), and all unspiked sewage samples (C). Bray-Curtis dissimilarities were calculated on Hellinger transformed data. The vegan function capscale was used to perform PCoA on the dissimilarity matrix. Variance explained by the two first axes are indicated. Sample replicates are connected with lines. For the results of the individual spiked samples as well as box plot visualizations, see Figures S4 and S5 in the supplemental material.

The pairwise dissimilarities for all samples were statistically assessed with ‘adonis’ and observed was a significant difference between samples (P<0.0001) (see Table S4 in the supplemental material). Since there were significant differences in group homogeneities, the analysis was supplemented by comparing dissimilarities with a Kruskal-Wallis test (see Table S5 in the supplemental material). The Kruskal-Wallis revealed also a significant difference between samples (P<0.0001), and that samples originating from the same input samples (P1, P2, S1, S2) were most similar to each other, respectively (see Table S5 in the supplemental material).

### A systematic effect on microbiome composition based on sample storage condition

As evident from the PCoAs, samples clustered in a systematic manner according to storage conditions for both, pig feces and sewage (Fig. 2B and C). The frozen samples (-80°C, -20°C) clustered together, and separately from the samples processed immediately (0h) and stored at higher temperatures (+5°C, +22°C). Of note, the microbiomes of the frozen samples remained stable over time (Fig. 2B, 2C, and see Fig. S5 in the supplemental material). In contrast, the microbiomes stored at positive °C differed dependent on the exact temperature they were stored at, i.e. the fridge (5°C) and room temperature (22°C). They also differed from samples processed right after collection (0h), and it appeared as the microbial communities of these samples (5°C, 22°C) changed along a time gradient in opposite direction of the frozen samples (-80°C, -20°C) (Fig. 2B, 2C, and see Fig. S5 in the supplemental material). However, the samples stored in the fridge (5°C) for 16h were in most cases relatively similar to the immediately processed samples (0h). A relatively large proportion of the variance was represented by the first axis, but more so for the pig feces samples as compared to the sewage samples. That storage had an effect on community composition was also confirmed in the ‘adonis’ test (P<0.0001) after subsetting to sample level (P1, P2, S1, S2). Model assumptions were in all cases fulfilled (P>0.05) (see Table S4 in the supplemental material).

The average dissimilarities of samples stored under different conditions relative to no storage (0h; immediate DNA isolation upon sample collection) revealed that the samples stored at 22°C for 64h had the largest dissimilarity relative to immediate DNA isolation, except for pig feces 1 (Table 1). In general, a large dissimilarity was observed for the frozen samples relative to 0h, but all frozen samples had a relatively small dissimilarity when compared with each other. Samples stored at 5°C for 16 hours mostly resembled samples that were processed immediately (Table 1, and see Table S4 in the supplemental material). These findings were in agreement with the PCoA.

**Table 1:** Average dissimilarities and dispersion (Standard error) of stored samples relative to samples that were not stored (i.e. DNA isolation immediately upon sample collection; 0h). The entries for 0h represented the variation among replicates.

|  | Pig feces 1 | Pig feces 2 | Sewage 1 | Sewage 2 |
| --- | --- | --- | --- | --- |
| 0h | 0,066 (0,003) | 0,081 (0,003) | 0,073 (0,003) | 0,057 (0,003) |
| 16h -80 °C | **0,159 (0,005)** | **0,116 (0,003)** | **0,103 (0,002)** | 0,077 (0,002) |
| 64h -80 °C | **0,152 (0,005)** | **0,113 (0,002)** | **0,121 (0,004)** | 0,076 (0,001) |
| 16h -20 °C | **0,141 (0,004)** | **0,124 (0,003)** | **0,109 (0,002)** | 0,082 (0,003) |
| 64h -20 °C | **0,139 (0,004)** | **0,126 (0,002)** | **0,109 (0,002)** | 0,088 (0,002) |
| 16h 5 °C | 0,073 (0,003) | 0,098 (0,004) | 0,089 (0,008) | 0,061 (0,003) |
| 64h 5 °C | 0,098 (0,002) | **0,137 (0,004)** | 0,091 (0,003) | 0,073 (0,001) |
| 16h 22 °C | 0,096 (0,002) | **0,130 (0,004)** | 0,087 (0,002) | 0,082 (0,002) |
| 64h 22 °C | **0,142 (0,002)** | **0,180 (0,003)** | **0,127 (0,002)** | **0,126 (0,001)** |

### Distinct taxa are driving the differences observed for different storage conditions

In order to determine the microorganisms that changed in abundance due to the different storage conditions, a constrained ordination was performed using canonical correspondence analysis (CCA) (Fig. 3A-D). A general trend across the four sample matrices was observed in that the majority of taxa related to the phyla *Firmicutes* and *Actinobacteria* were associated with samples stored at positive degree C (5°C, 22°C). In contrast, *Bacteroidetes* and *Proteobacteria* were more associated with the frozen samples (-80°C, -20°C) (Fig. 3A-D, see also Fig. S6 in the supplemental material).

**Figure 3:**
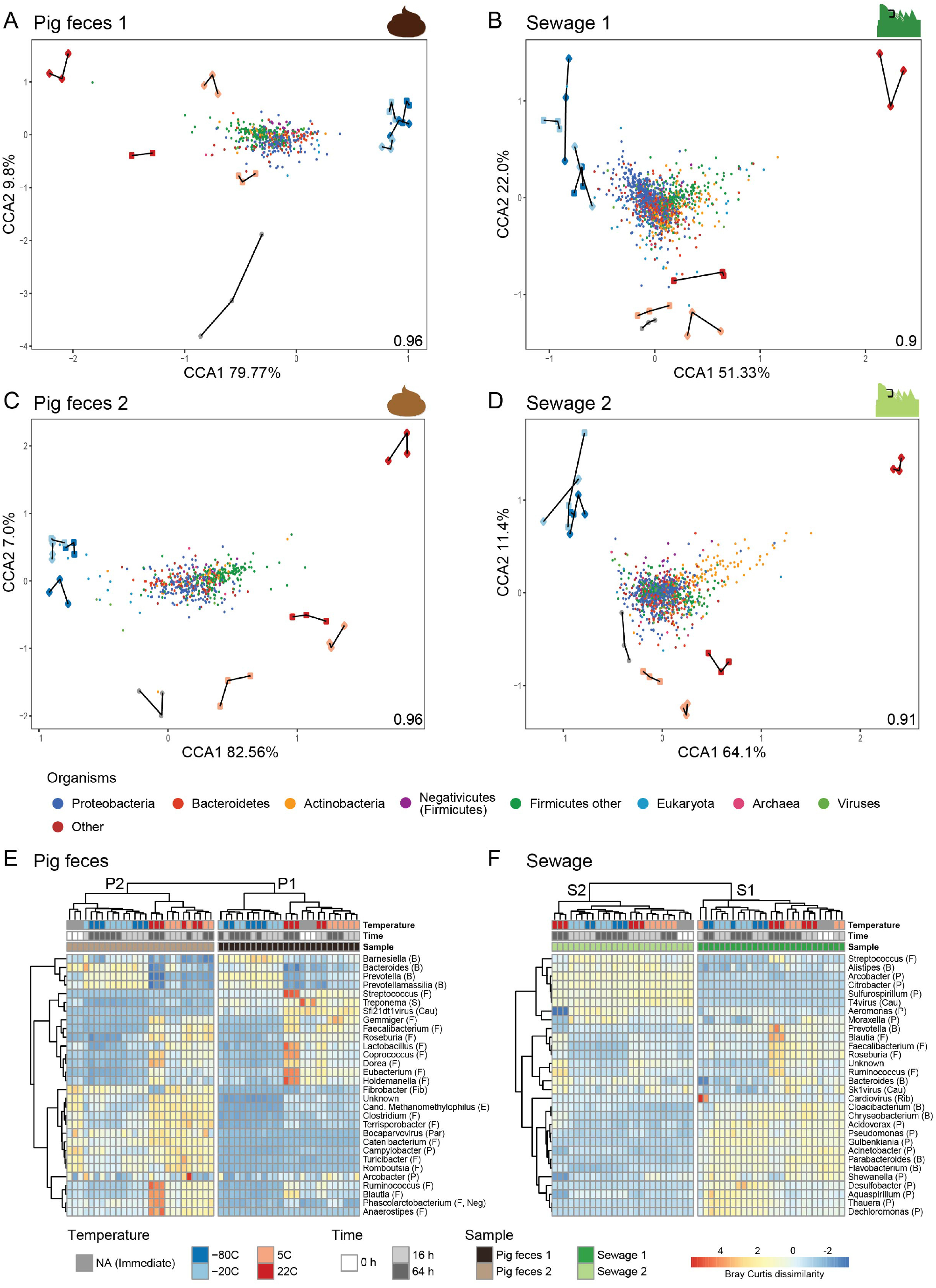
Overview of community composition in relation to storage conditions. (A-D) Ordination plots of canonical correspondence analyses (CCA) of the individual unspiked samples (P1, P2, S1, and S2) visualizing different groups of organisms (coloured points) constrained by the storage conditions (coloured shapes with replicates connected with lines). The inertia constrained by the explanatory variables is indicated in the lower right corner of the CCA plot, respectively. Heatmaps displaying the 30 most abundant genera in unspiked pig feces (E) and unspiked sewage (F) using Bray-Curtis dissimilarity matrix calculated on Hellinger transformed data. Complete-linkage clustering was used for both genera and samples. The clustering of genera was performed on Pearson correlation coefficients and samples. For the heatmap visualization genera were standardized into zero mean and unit variance. Phyla are indicated as B: Bacteroidetes, F: Firmicutes, S: Spirochetes, Cau: Caudovirales, Fib: Fibrobacteres, E: Euryarchaeota, Par: Parvoviridae, P: Proteobacteria, Neg: Negativicutes, Rib: Riboviria. For the results of the individual spiked samples, see Figure S6 in the supplemental material.

To elucidate the patterns in community structure in more detail, a cluster analysis of pig feces and sewage samples was performed at genus level and visualized in heatmaps (Fig. 3E and F). The two pig feces and two sewage samples clustered separately, respectively, supporting the observations from the PCoA. In agreement with the CCA, the cluster analysis of the pig feces samples also revealed that *Firmicutes* such as *Faecalibacterium*, *Coprococcus*, *Eubacterium*, *Ruminococcus*, and *Blautia,* were more abundant in samples stored at positive degree C (5°C, 22°C), compared to the other storage conditions (Fig. 3A, C and E). Furthermore, the observation for the pig feces samples that *Bacteroidetes* and *Proteobacteria* were more abundant in frozen samples was also confirmed in part by the cluster analysis in which genera such as *Barnesiella*, *Bacteroides*, *Prevotella* and *Prevotellamassilia* were observed in higher abundance compared to the other samples (Fig. 3A, C and E). The patterns related to Firmicutes, Bacteroidetes, and Proteobacteria in the CCA compared to the cluster analysis were less clear for the sewage samples (Fig. 3B, D and F). Some

Firmicutes genera appeared indeed to be present in higher abundance in samples stored at positive degree C, such as *Blautia*, *Faecalibacterium*, *Roseburia*, and *Ruminococcus*, compare to the other storage conditions (Fig. 3F). Overall for the sewage samples, it was more difficult to reveal which genera were driving the differences between storage conditions using this type of analysis.

In the CCA, we observed that Firmicutes and Actinobacteria abundances were associated with storage at positive degree Celsius, and Bacteroidetes and Proteobacteria with frozen samples (Fig. 3). We assessed this finding further using MA plots with log2 fold changes from a differential abundance analysis using DESeq. We focussed on storage conditions for which the highest dissimilarities were observed among each other, i.e. samples stored at 22°C, 5°C and -80 °C for 64 hours, respectively, compared to immediate DNA isolation (see Fig. S7-9 in the supplemental material). The most striking patterns were observed when comparing samples stored at 22°C for 64 h compared to samples undergoing immediate DNA isolation (see Fig. S7 in the supplemental material). *Firmicutes* and *Actinobacteria* were observed at higher abundance upon storage at 22°C for 64 h, and a corresponding lower abundance was observed for *Bacteroidetes* and *Proteobacteria* compared to samples processed immediately (see Fig. S7 in the supplemental material). The *Negativicutes* as part of *Firmicutes* tended to exhibit similar abundance patterns as *Firmicutes*. The effect also seemed independent of abundance with an expected larger variance of the lower abundant genera (see Fig. S7 in the supplemental material). The same patterns were observed when comparing immediate DNA isolation with the samples stored at 5°C for 64 h (see Fig. S8 in the supplemental material). Comparing immediate DNA isolation with samples stored at -80°C for 64 h revealed that the *Negativicutes* within *Firmicutes* were almost exclusively more abundant in the frozen samples, and no clear patterns at phylum level were otherwise observed (see Fig. S9 in the supplemental material).

### Distinct genera are not always exhibiting the same abundance patterns across sample types

To quantify the effect of storage conditions on microbial community composition, we tested for the differential abundance of genera in pairwise comparisons using DESeq analyses that did not take the compositionality of the data into account (Fig. 4 and dataset 1 available from https://doi.org/10.6084/m9.figshare.12010983). *Prevotella* (class Bacteroidia), the most abundant genus in pig feces, was for both samples (P1, P2) more abundant in frozen samples compared to the samples undergoing immediate DNA isolation. At positive degree Celsius, *Prevotella* seemed to decrease in relation to temperature and time, i.e. *Prevotella* abundance decreased the higher the storage temperature was, and the more time passed (Fig. 4 and dataset 1 available from https://doi.org/10.6084/m9.figshare.12010983). A decrease in the most abundant genus in pig feces might also explain the observed increase in evenness (Peilou’s evenness) in pig feces (Fig. 1B). *Faecalibacterium* (class Clostridia) in pig feces exhibited the opposite pattern compared to *Prevotella*, and was detected in both samples (P1, P2) at a higher abundance with increasing temperature and time, and lower abundance in frozen samples, compared to the samples processed immediately. *Treponema* (class Spirochaetes) was observed at a higher abundance in the samples processed immediately upon sample collection for both pig feces samples (P1, P2), compared to the stored samples, independent of temperature and time. The opposite was observed for *Phascolarctobacterium* (class Negativicutes), for which a higher abundance was observed for stored samples compared to the samples processed immediately.

**Figure 4:**
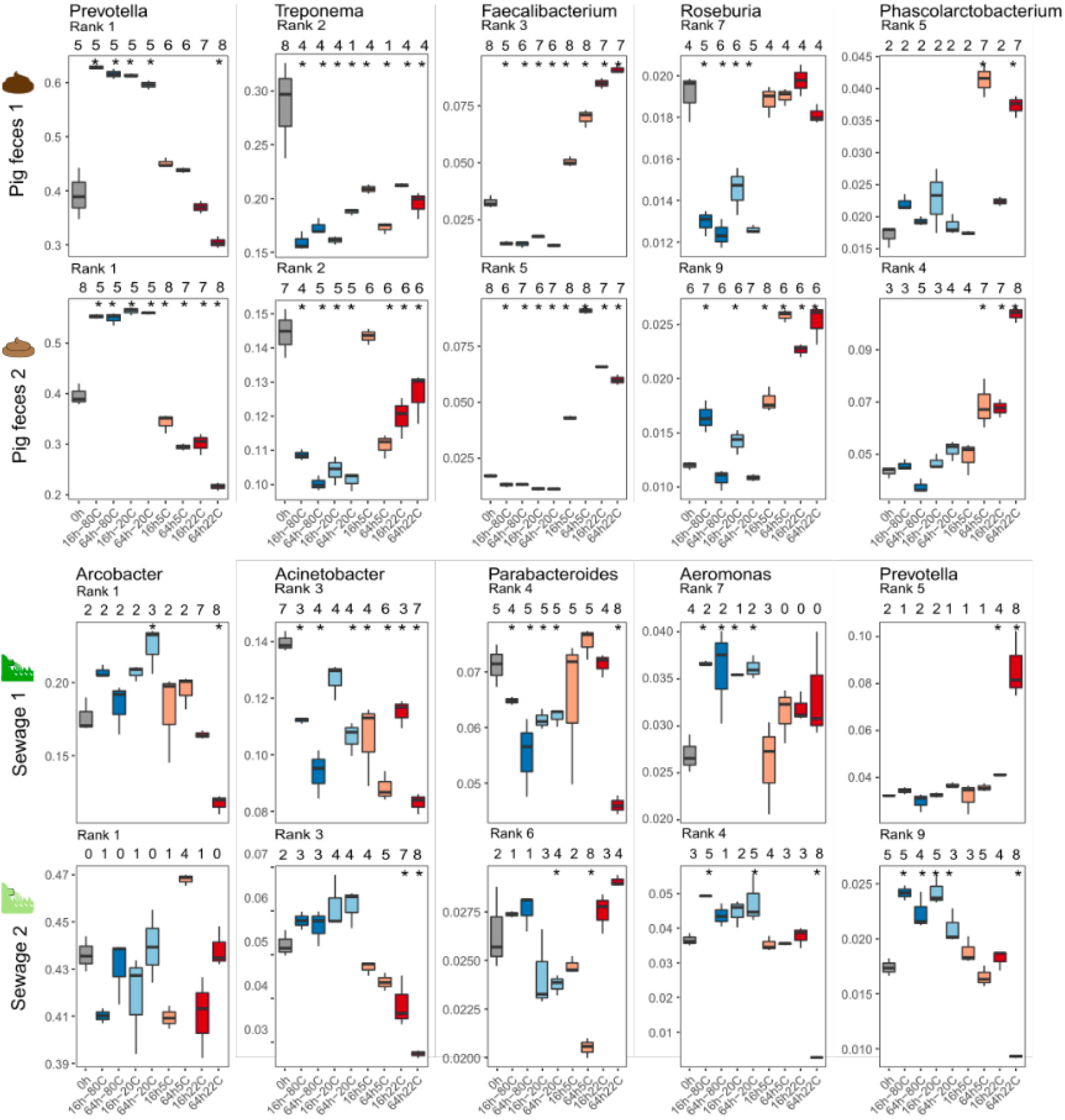
Boxplots displaying the relative abundance of selected highly abundant bacteria in unspiked pig feces (P1, P2) and sewage (S1, S2). The abundance rank is indicated for each genus in the specific sample. Differential abundance between the storage conditions was tested pairwise using DESeq2, and the number of times a storage condition was significantly different (p<0.01) from another is indicated above the boxplots. The asterisks indicate storage conditions that were significantly different from immediate DNA isolation at 0h.

The two sewage samples (S1, S2), in contrast to the pig feces samples, exhibited less concordant genus abundance patterns between each other, respectively, and to the pig feces samples (Fig. 4). However, as also observed for the two pig feces samples, *Prevotella* in Sewage 2 was detected at the highest abundance in frozen samples compared to the other storage conditions (Fig. 4). Overall, some generalizable genera abundance patterns were observed. However, some differences were observed between samples from the same environment (e.g. S1 vs S2) and different environments (pig feces vs sewage), indicating that the effect of storage on specific genera is not entirely generalizable across samples from the same and different environments (Fig. 4 and dataset 1 available from https://doi.org/10.6084/m9.figshare.12010983).

### Effect of storage conditions on mock community

Duplicate sets of pig feces and sewages samples were spiked with a mock community composed of 8 different microorganisms and stored at the different temperatures for different periods of times (Fig. 1A, and see Tables S2 and S3 in the supplemental material). Overall, the abundance patterns of these microorganisms under the examined storage conditions appeared individualized for P1, P2, S1, and S2, with few exceptions (see Fig. S10A in the supplemental material). A generalizable effect was observed across all four samples for *Cryptosporidium,* resulting in higher relative abundances in samples that were frozen (-20°C, -80°C) compared to the samples that were processed immediately (0h). The effect was most pronounced for the samples stored at -80°C (see Fig. S10A in the supplemental material). In addition, for all samples, a decrease in *Staphylococcus* and *Fusobacterium* abundances was observed compared to the samples that were processed immediately (0h). This effect was most pronounced for the sewage samples and independent of storage temperature and time. Furthermore, an increase of *Saccharomyces* was observed for several samples, for example P1, S1 and S2 at 22°C, while a decrease was observed for example for S1 and S2 at both, -80°C and -22°C (see Fig. S10A in the supplemental material). However, the these taxa did not exhibit the same abundance patterns in spiked and unspiked samples, even though the six bacterial genera were detected in many of the unspiked samples as part of the native microbial community (see Fig. S10A and B in the supplemental material). For example, in the unspiked samples, *Salmonella* in P2 exhibited a decrease in abundance at almost all storage conditions compared to the samples processed immediately. This was not observed for the respective P2 samples spiked with additional cells of *Salmonella* added to the pig feces microbial community (see Fig. S10A and B in the supplemental material).

When the abundance of the mock community strains residing in the pig and sewage samples is analysed in relative abundance to each other, it appears that the abundance of Gram-negative bacteria is higher compared with Gram-positive bacteria and the eukaryotes (*Cryptosporidium, Saccharomyces)* (see Fig. S10C in the supplemental material). Nevertheless, also here one can observe a decrease of *Fusobacterium* and *Staphylococcus* abundances under certain storage conditions as described above (see Fig. S10 in the supplemental material). When the abundance of the mock community strains prior to spiking is analysed based on cells counts and CFU, it appeared that the abundance of Gram-positive bacteria is higher compared with Gram-negative bacteria and the eukaryotes (see Fig. S10C in the supplemental material). When DNA from the individual mock community strains was isolated using the same DNA extraction protocol that was optimized and used for complex microbiome samples, more DNA was obtained from the Gram-negative bacteria compared to Gram-positive bacteria and the eukaryotes (see Fig. S10C in the supplemental material). However, the concurrent analysis of both spiked and unspiked sets of samples throughout this study revealed that overall the same observations were obtained in regards to microbial community and antimicrobial resistome patterns for both sets of samples (e.g. compare Fig.1B and S3, S4, S5, 5 and S13), also providing an additional layer of technical control in this study.

**Figure 5:**
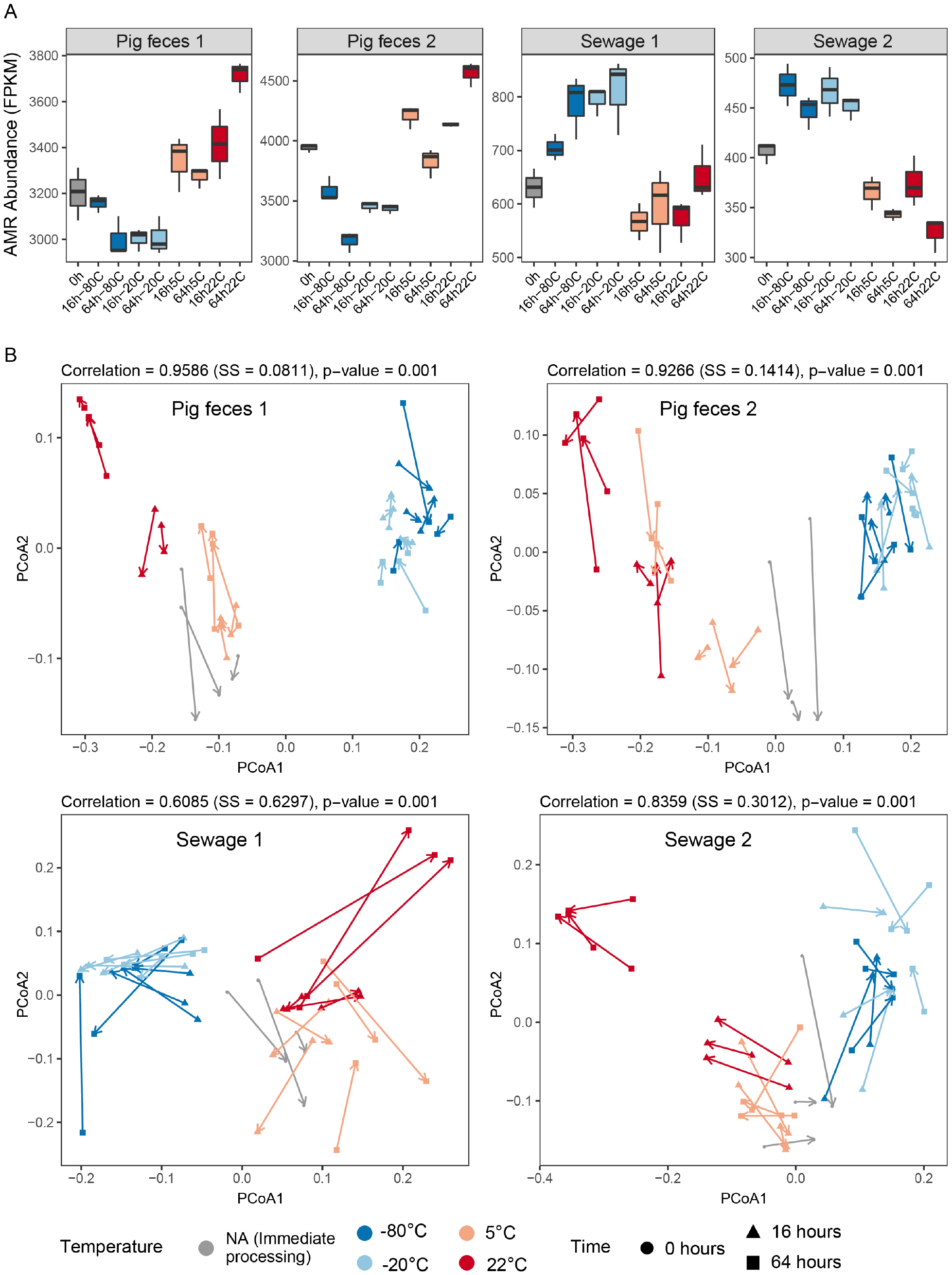
The effect of sample storage conditions on the resistome. A) Boxplots displaying total antimicrobial resistance (AMR) abundance in the unspiked samples (P1, P2, S1, S2) measured in FPKM relative to the total number of bacterial reads. B) Procrustes rotation comparing the resistome and taxonomic dissimilarities in the four unspiked samples. For the results of the spiked samples, see Figure S13 in the supplemental material.

### Effect of freeze-thaw cycles on microbial community composition

In order to assess the effect of repeated freeze-thaw cycles on inferred microbial community composition, separate aliquots of pig feces P1 and sewage S1 were frozen at -20°C and -80°C at 0h, thaw and refrozen after 16h, and thaw and frozen again after 40h (2 freeze-thaw cycles), 64h (3 freeze-thaw cycles), and 88h (4 freeze-thaw cycles) (Fig. 1A, and see Table S1 in the supplemental material). We examined the inferred microbial community composition in PCoAs, and it appeared as that the communities changed on a gradient according to an increasing number of freeze-thaw cycles, in particular for pig feces P1 (see Fig. S11A in the supplemental material). To reveal the microorganisms that changed in abundance due to the repeated freeze-thaw cycles, CCAs were performed. For both, pig feces and sewage, some eukaryotic genera as well as Actinobacteria and Firmicutes were more associated with samples undergoing increasing numbers of freeze-thaw cycles (see Fig. S11B in the supplemental material). In regard to the eukaryotic genera, we for example observed in the spiked samples an increasing abundance of *Cryptosporidium* with increasing numbers of freeze-thaw cycles (see Fig. S11C in the supplemental material). In contrast, Proteobacteria and Bacteroidetes appeared to be more associated with low numbers of freeze-thaw cycles (see Fig. S11B in the supplemental material).

### A systematic effect on antimicrobial resistomes based on sample storage condition

To examine whether antimicrobial resistance (AMR) patterns would change in response to the different storage conditions, we mapped the reads from all samples against the ResFinder database. A rarefaction analysis revealed that the majority of acquired AMR genes discoverable under these conditions was observed for the pig feces samples, revealed through a flattening rarefaction curve (see Fig. S12A in the supplemental material). This was not the case for the sewage samples that exhibited a higher AMR gene richness and lower total number of mapped reads compared to the pig feces samples (see Fig. S12A in the supplemental material). Most AMR genes in pig feces represented genes conferring resistance to tetracycline (average abundance = 69.0%), and in sewage against macrolides (average abundance = 31.7%) (see Fig. S12B in the supplemental material). The second and third most abundant AMR classes in sewage represented AMR genes against tetracyclines (average abundance = 24.8%) and beta-lactams (average abundance = 23.6%) (see Fig. S12B in the supplemental material).

Overall, the total observed abundance of AMR genes appeared to be depending on the storage conditions, and opposite patterns were observed for pig feces and sewage samples (Fig. 5A, and see Fig. S13A in the supplemental material). The total AMR abundance was lowest in pig feces (P1, P2) when samples were frozen (-20°C, -80°C) and in sewage (S1, S2) when stored at positive degree C (22°C, 5°C). In contrast, total AMR abundance was highest in pig feces (S1, S2) when stored at positive degree C (22°C, 5°C), and in sewage (S1, S2) when frozen (-20°C, -80°C) (Fig. 5A, and see Fig. S13A in the supplemental material).

Comparing the resistome with the taxonomic patterns using procrustes analysis revealed that in all cases (P1, P2, S1, S2) they correlated significantly (P<0.001), respectively (Fig. 5B and see Fig. S13B in the supplemental material). Storage had a systematic effect on the inferred taxonomic and antimicrobial resistome-pattern, particularly the frozen samples that clustered together. Even though a clustering of samples stored at positive °C was also observed, the resistome-based patterns were less pronounced compared to the taxonomic-based patterns (Fig. 5B and see Fig. S11B in the supplemental material).

### AMR classes exhibit distinct abundance patterns under different storage conditions

An analysis of the most abundant AMR classes revealed different effects of storage on the resistome across the sample types (pig feces, sewage). For pig feces (P1, P2), tetracycline-resistance genes were detected at a higher abundance in samples stored at positive degree C (22°C, 5°C) and lower in samples when frozen (-20°C, -80°C) compared to samples processed immediately (Fig. 6 and see Fig. S14 in the supplemental material). A similar pattern was observed for aminoglycoside- and lincosamide-associated resistance genes. In contrast, macrolide- and beta-lactam associated genes were detected at a higher abundance in samples stored frozen (-20°C, -80°C) compared to samples processed immediately and stored at positive degree (22°C, 5°C) (Fig. 6 and see Fig. S14 in the supplemental material). In sewage (S1, S2), the pattern for the top 4 most abundant AMR classes (macrolide, tetracyclin, beta-lactam, aminoglycoside) was similar in that in most cases AMR class abundance was highest when samples were stored frozen (-20°C, -80°C) (Fig. 6 and see Fig.S14 in the supplemental material). Macrolide and beta-lactam resistance genes abundance patterns were similar as observed in pig feces samples, i.e. AMR gene abundance was higher in frozen samples (-20°C, -80°C) compared to samples stored at positive °C (22°C, 5°C).

**Figure 6:**
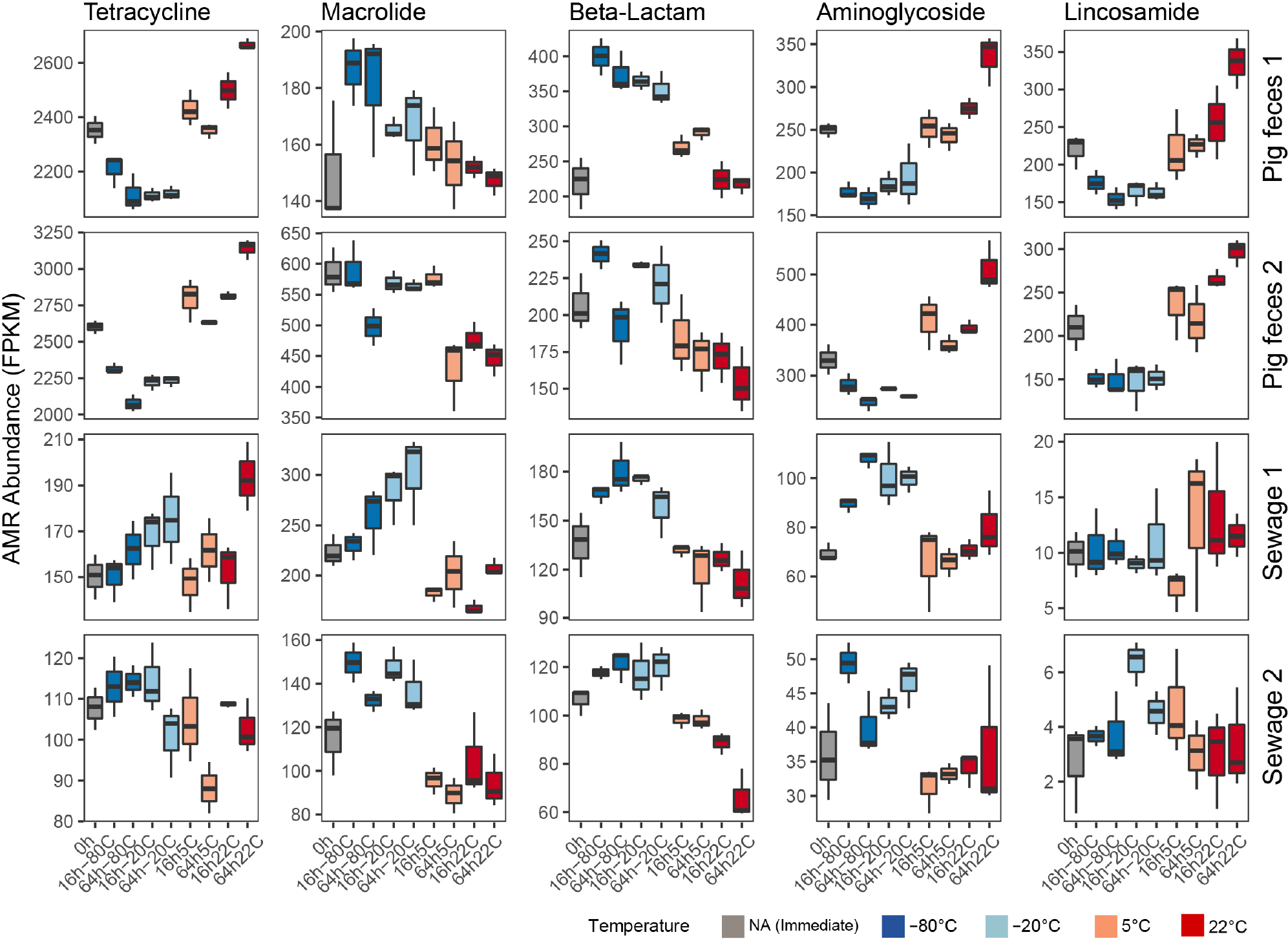
The effect of storage on the abundance of antimicrobial resistance classes. Boxplots displaying total antimicrobial resistance (AMR) abundance for the most abundant antimicrobial resistance classes in the unspiked samples (P1, P2, S1, S2). The abundance was measured in FPKM relative to the total number of bacterial reads. For the results of the spiked samples, see Figure S14 in the supplemental material.

## Discussion

Metagenomics is a powerful technology to obtain insight about the taxonomic and functional composition of microbial communities. Standardization and harmonization of the laboratory procedures for metagenomics-based analyses are important, in particularly for long-term studies that aim at investigating variations in microbial community composition over time, and it would also facilitate meta-analyses involving data from different studies. Knowledge about the impact of different sample processing strategies would furthermore allow designing more precise algorithms for the identification and correction of batch effects. In general, it is critical to know which effect different sample processing steps have in skewing the true microbial community composition to facilitate improving laboratory protocols and more rigorous interpretations of metagenomics-based analyses.

We here focussed on the impact of one of the key initial steps in metagenomics, namely the storage conditions for biospecimens. This step can be difficult to standardize, and we chose to investigate different real-world scenarios. For example, in regard to storage duration times we investigated immediate sample processing (0 hours), as well as storage for 16 hours and 64 hours resembling situations where samples are received in the afternoon and sample processing can first initiated the following day or after a weekend, respectively. We also aimed at examining situations where samples have to be stored for extended periods of times (4, 8, 12 months). The storage temperatures were chosen to reflect common ways to store samples such as in the fridge (5°C), freezer (-20°C) and deep freezer (-80°C), but also storage temperatures that might be experienced were cooling and freezing is not an option (e.g. 22°C), such as in field studies. In addition, we examined the effect of freeze-thaw cycles, a scenario where one has to repeatedly retrieve aliquots from the biospecimen stored in the freezer, for example for additional rounds of DNA isolation, or other types of analysis such as metabolomics, and where not the entire specimen is kept in the frozen state.

Overall, we found that the four investigated biospecimens could still be distinguished, independent of the effect introduced by the different storage conditions, as they clustered by sample origin and not based on storage condition. This is in support of previous findings (11). However, we anticipate that it would not necessary always be possible to trace back samples to their origin using standard metagenomics analyses if one is examining a larger set of samples of the same type (e.g. pig feces). Therefore, it is advised to store all samples in the same way, and ideally under frozen conditions (-20°C or -80°C) under which we observed the least changes in microbial community composition over time.

We found that storage had a systematic effect on the inferred taxonomical microbiome composition. For example, frozen samples (-20°C or -80°C) clustered together and were more similar to each other than samples stored at positive degree Celsius (5°C and 22°C). It should be noted that while freezing the samples changed the detected microbiome compared to immediately processing the samples, the results were stable over time. Previous 16S rRNA-based studies on human feces have found different results in regard to the effect of different sample storage conditions on alpha and beta diversity, as well as taxa abundances changes (13, 15, 16).

We also observed that the storage conditions had a systematic effect on the antimicrobial resistome that correlated with the taxonomic analysis. For all samples stored at positive degree C (5°C, 22°C) we observed a higher abundance of taxa related to the phyla *Firmicutes* and *Actinobacteria*, and a lower abundance of taxa related to *Bacteroidetes* and *Proteobacteria* which were more associated with the frozen samples (-80°C, -20°C). This may be explained by an increased release of DNA from some Gram-negative bacteria such as *Bacteroidetes* and *Proteobacteria* upon freezing as compared to samples that lack this step (e.g. samples processed immediately). Of note, while the most abundant genera exhibited abundance changes similarly in both pig feces samples (e.g. *Prevotella, Treponema, Faecalibacterium*), this was not the case for the two sewage samples. This could be impacted by a higher degree in differences in chemical compositions, physical properties, and water content in the sewage samples compared to the pig feces (2, 17–19). This may also be supported by the observation that both alpha and beta diversity were more similar between the two pig feces samples as compared to the two sewage samples, respectively.

We spiked a subset of all samples with a well-characterized mock community and examined abundance changes of these in response to the different storage conditions. Interestingly, the members of this mock community did not seem to respond to the storage conditions in a manner similar to the close relatives of the native community, even though the six bacterial genera were present as well in many of the unspiked samples as part of the native microbial community. We cultivated the mock community members on plates in the laboratory. Hence, they might not exhibit the same physiological state as the native members, not be able to rapidly accommodate to the new conditions and respond differently to the DNA isolation procedure (2). Differences between the theoretical and observed abundances of spiked-in bacteria have been previously observed (20–24). In contrast to the bacteria and fungus, the *Cryptosporidium* oocysts are likely to behave similar in this setting compared to natural settings due to the dormant nature of this stage. The findings of an increase in *Cryptosporidium* under freezing conditions are therefore of relevance. It should though be mentioned that under clinical settings a sample may additionally contain other stages of *Cryptosporidium* for which improved lysis might not require freezing.

In the sub-experiment, involving repeated freeze-thaw cycles, we observed an increase in the abundance of eukaryotes. For studies aimed at detecting microbial eukaryotes, freezing the biospecimens seems therefore to have a positive impact on sensitivity. For instance, the clinically relevant *Cryptosporidium parvum* that was part of the mock community had a relatively higher abundance in samples that underwent 3-4 freeze-thaw cycles as compared to 2 freeze-thaw cycles. Furthermore, an increasing abundance of Actinobacteria and Firmicutes was observed upon increasing the number of freeze-thaw cycles, suggesting that this procedure facilitated a disruption of more rigid cell walls. However, repeated freeze-thaw cycles may also contribute to a degradation of DNA that is more easily released from Gram-negative bacteria, contributing to a decreasing abundance observed for these taxa. This is supported by the finding that a higher abundance of Bacteroides and Proteobacteria was more associated with samples undergoing 2 freeze-thaw cycles as compared to 3-4 freeze-thaw cycles.

Overall, we recommend that all samples are stored in the freezer (-20°C or -80°C) prior to DNA isolation. While microbial community composition remained stable under these conditions over shorter periods of time, it remains to be investigated more thoroughly whether this would also be the case for extended periods of time, as samples within a study might be stored for different lengths of time. The strategy of storing samples in the freezer could be an advantage as generally most samples are stored frozen, facilitating better comparability between studies. If it is not possible to store samples in the freezer, it would be good to perform DNA isolation immediately upon arrival at the lab if feasible for all samples in a study. If this is not possible, we recommend to store samples in the fridge and process them the same day or the following day for all samples. It could also be possible to store samples using preservation reagents (12, 13, 25); however, these interventions might interfere with other investigations that are also to be conducted on the same biospecimen (e.g. metabolomics, proteomics, transcriptomics, cultivation). Within a project, standardization of storage conditions is key for generating comparable data. We strongly recommend that more details regarding sample storage and processing conditions are provided as part of the metadata submitted together with the DNA sequencing data to public repositories than is currently done (26).

## Materials and Methods

### Microbiome samples

Two pig feces and two domestic sewage samples were collected for this study. The pig feces samples were obtained from two individual animals right after defecation at two different conventional pig production farms in Denmark on separate occasions and were transported within 3 hours to the laboratory in a cooling box. The unprocessed domestic sewage was collected at a local waste water treatment facility (Lyngby-Taarbaek forsyningen) on separate occasions as 20 L inlet water in 5-liter sterilized plastic bottles, transported in cooling boxes to the laboratory within 20 minutes, and placed into a fridge at 5°C. Sedimentation of the sewage was commenced immediately using 50 ml Falcon tubes and two centrifuges (Eppendorf 5810R, Hamburg, Germany. Program: 10 min at 10,000 g). Sedimentation of the entire sewage sample was completed within 12 hours.

Each individual pig feces sample and sewage pellet were thoroughly homogenized in a 50 mL falcon tube with a sterile wooden spatula and distributed into two large aliquots in 50 mL falcon tubes, respectively. One aliquot for each of the four original microbiome samples, respectively, was spiked with a freshly prepared mock community (see below), and all samples were homogenized with a sterile wooden spatula to take into account that the mixing might have an altering effect on community composition. Small aliquots for each sample storage condition were prepared in Eppendorf tubes, and either processed immediately (storage for 0h) or stored at a certain temperature (-80°C, -20°C, 5°C, or 22°C) for a specific period of time (16h, 64h, 4 mo, 8 mo, 12 months) (described in detail below). Additional aliquots underwent 2-4 freeze thaw cycles at different time points (40h, 64h, 88h) (described in detail below).

### Mock community

In parallel to the microbiome sample collection and initial processing, a mock community was prepared at the four individual occasions. The mock community consisted of eight different microorganisms: *Propionibacterium freudenreichii* DSM20271, *Bacteroides fragilis* NCTC9343, *Staphylococcus aureus* NCTC8325, *Fusobacterium nucleatum* ATCC25586, *Escherichia coli* ATCC25922, *Salmonella enterica* ATCC14028S, *Cryptosporidium parvum* Iowa II isolate, and *Saccharomyces cerevisiae* S288C, representing two different domains of life (Bacteria, Eukarya) and seven phyla. The bacterial domain was represented by five phyla (*Actinobacteria*, *Bacteroidetes*, *Firmicutes*, *Fusobacteria*, *Proteobacteria*), which differed in terms of cellular morphology (cocci, rods, and fusiform rods), cell wall structure (Gram-positive, Gram-negative) as well as oxygen requirements. The eukaryotic domain was represented by two different phyla (*Apicomplexa*, *Ascomycota*). The particular microorganisms were also chosen because their whole genome sequence was publicly available. Additional information about the mock community microorganisms including cultivation conditions is provided (see Table S2 in the supplemental material). The microorganisms were transferred to an Eppendorf tube (1.5 ml) with an inoculation loop and resuspended in sterile PBS (Gibco, Paisley, UK) with a table-top vortex mixer. Enumeration of the bacteria and the fungus was performed using a Petroff counting chamber under a light microscope, counting two diagonal corners on two separately prepared slides. Organisms cultured aerobically (*S. aureus*, *E. coli*, *S. enterica*, and *S. cerevisiae*) were processed first. Raw cell counts and volumes of resuspended cells that were used for the spiking of fecal and sewage samples are provided (see Table S3 in the supplemental material). The *C. parvum* oocysts were obtained from Waterborne Inc. in PBS amounting to 1.2*10^8 number of cells, as determined by FACS by the provider. The oocysts were washed 5x by resuspending them in 10 ml of PBS and sedimenting them using a centrifuge (Eppendorf 5810R, Hamburg Germany, 1000 g, 10 min.). The mock community was mixed, leading to 10^9 (for P1: 5*10^8) cells/mg Gram-positive bacteria, 10^8 cells/mg Gram-negative bacteria, 2*10^7 cells/mg *S. cerevisiae* and 2*10^6 cells/mg *C. parvum*, respectively, in the pig feces and pelleted sewage samples, respectively (see Table S3 in the supplemental material). Colony forming units (CFU) counts were obtained subsequently to estimate the number of live cells. CFU were determined for selected organisms and was depending on the microbiome sample that was processed (see Table S3 in the supplemental material).

### Sample storage conditions

The effect of different sample storage conditions was investigated at four temperatures (deep freezing: -80°C, freezing: -20°C, refrigerator 5°C, and room temperature 22°C) and six storage duration times representing relevant situations in microbiome studies (immediate DNA isolation: 0 hours; sample storage overnight: 16 hours; and sample storage over a weekend: 64 hours; and longer-term sample storage: 4 mo, 8 mo, 12 months). To assess different sample matrices and different samples of the same matrix, two pig feces samples (P1 & P2) and two sewage samples (S1 and S2) were included (for long-term storage: P1 and S1, only). The technical variability was accounted for by processing all sub-samples in triplicates from the step of DNA isolation.

In addition, given that in microbiome studies sample aliquots may be repeatedly retrieved from the main sample stored in a freezer, we aimed at simulating this in a sub-experiment by a series of freeze-thaw cycles. Aliquots were placed in the freezer (-20°C) and deep-freezer (-80°C), thawn and refrozen after 40h (2 freeze-thaw cycles), 64h (3 freeze-thaw cycles), and 88h (4 freeze-thaw cycles). An overview of all 343 samples, including controls, is provided in Table S1 in the supplemental material.

### DNA isolation and metagenomics shotgun sequencing

DNA isolation was performed according to a modified QIAamp Fast DNA Stool Mini Kit (Qiagen) protocol that included a bead beating step with MoBio garnet beads, and additional steps optimized for pig feces and sewage (https://doi.org/10.6084/m9.figshare.3475406) (2). Using the same DNA isolation procedure, DNA was also isolated from aliquots of each individual strain of the mock community. A DNA isolation blank control was included at each round of DNA isolation. Positive controls consisted of the pure mock community, prepared at the four individual occasions and they were processed in triplicates together with the other samples at the 0 hour time points, respectively. DNA concentrations were measured with the Qubit dsDNA High Sensitivity (HS) assay kit on a Qubit 2.0 fluorometer (Invitrogen, Carlsbad, CA) before storing the DNA at -20°C.

Metagenomics shotgun sequencing was for the majority of samples (i.e. 287 samples) performed on an Illumina HiSeq 4000 (2x150 cycles, paired end) at Oklahoma Medical Research Foundation (USA). DNA was processed for sequencing, involving mechanical fragmentation (Covaris E220evolution, aimed insert size=350bp), and followed by amplification-free library preparation using NEXTflex PCR Free DNA prep kit (Bioo Scientific) according to standard manufacturer protocols. The remaining samples (i.e. 56 samples) associated with long-term storage (4, 8, 12 months) of P1 and S1 were sequenced on an Illumina HiSeq 4000 (2x150 cycles, paired end) at Admera Health, New Jersey (USA). DNA was processed for sequencing, involving mechanical fragmentation (Covaris E220evolution, aimed insert size=350bp), followed by amplification-free library preparation using KAPA library preparation kit (Roche) according to standard manufacturer protocols. Samples were distributed randomly across flow cell lanes. Labelling issues occurred for four samples resulting in the swapping of these samples. It was possible to backtrack the samples that were mislabelled based on labbook entries, DNA concentration measurements performed at both, the research laboratory and sequencing provider, and using principal coordinates analysis of Bray-Curtis distances performed corresponding to the PCoA analysis described in detail below. These four samples, of which one was a DNA isolation blank control, were excluded from further analysis to assure mislabelling did not have an effect on the interpretation of results. Furthermore, one sample was excluded due to low sequencing output.

### Sequencing data analysis

Demultiplexed raw reads were processed with BBduk2 as part of the BBTools package (http://jgi.doe.gov/data-and-tools/bbtools/) to remove low-quality bases from reads (trimq=20), remove reads shorter than 50 bp and adapter sequences. Mapping of processed paired-end (PE) reads was performed with MGmapper (27) that incorporates a Burrows-Wheeler aligner (BWA-mem). Reads were mapped in bestmode against 11 databases (human, bacteria, bacteria_draft, fungi, archaea, virus, parasites_vertebrates, parasites_other, HumanMicrobiome, common_animals, and common_plants) containing genome sequences that were downloaded from NCBI (see Table S6 in the supplemental material) with a 0.8 fraction of matches+mismatches relative to the full length of the read (27). Details about the number of raw reads (average: 14.4 million PE reads, range: 2.9-41.0 million PE reads per sample), number of reads that passed the quality filtering (average: 12.4 million PE reads, range: 2.5-35.9 million PE reads), and mapped reads (average: 2.2 million PE reads, range: 0.65-7.1 million PE reads) are provided in (see Table S1 in the supplemental material). The genome assembly sizes where used for the normalization according to genome size. In addition, increased hit counts to specific contigs were adjusted as implemented previously (6).

The taxonomic-based analyses were performed with reads mapping to genomes of all bacteria, archaea, viruses, fungi and Cryptosporidium, and analyses were performed by aggregating counts to the level of genera. The mapping to functional databases (ResFinder, Virulencefactors, Virulence) was performed in “fullmode” (allowing reads to be assigned to multiple databases), and the same mapping criteria as described above had to be fulfilled. ResFinder was the only database that was used for further analysis and includes acquired resistance genes and chromosomal point mutations (28). Resistance genes were aggregated with respect to resistance against different antimicrobial classes.

### Statistical analysis

#### Pre-processing

The read count table was first processed by dividing all counts with two, since mapping was performed as proper pairs (mapping of a forward and reverse read), this is referred to as the raw count table (see item A at https://doi.org/10.6084/m9.figshare.12010989). The raw count table was normalized according to genome size for each microorganism, followed by total sum scaling. The resulting table is referred to as the count table (see item B and C at https://doi.org/10.6084/m9.figshare.12010989). The microorganisms listed in the tables were aggregated to genus level. For the antimicrobial resistome analysis, raw read counts are provided for all genes divided by two (see item D at https://doi.org/10.6084/m9.figshare.12010989) and normalized according to the total bacterial reads per sample (mapping to the databases: bacteria, bacteria_draft and HumanMicrobiome) as fragments per kilo base reference per million bacterial fragments (FPKM) (see item E at https://doi.org/10.6084/m9.figshare.12010989). The FPKM table was also aggregated to resistance against antimicrobial classes (see item F at https://doi.org/10.6084/m9.figshare.12010989), and the class-level information for each AMR-genes are available as item G at https://doi.org/10.6084/m9.figshare.12010989.

All statistical analysis and visualization of data was performed in R (Version 3.4.4, R Foundation for Statistical Computing, Vienna, Austria) (29). The multivariate community ecology analysis was performed with vegan version 2.5.2 (30) and the differential abundance analysis was performed with DESeq2 version 1.18.1 (31). Visualisation of data was performed with ggplot2 version 2.3.0 (32) and pheatmap version 1.0.10. All analyses were performed separately for both spiked and unspiked samples. Final modifications to figures were made in Inkscape or Illustrator.

A high degree of concordance was observed between the unspiked and spiked samples. The majority of data presented in the main text are based on the unspiked samples. The same analyses of the spiked samples are presented in the supplementary material.

#### Alpha-diversity

Alpha diversity was calculated based on the raw count table estimating richness (Chao1), evenness (Pielou’s) and diversity (Simpson) using the diversity function in vegan. Boxplots were created for each sample and according to the different storage conditions.

#### Dissimilarity calculations

A dissimilarity matrix was generated based on Bray-Curtis dissimilarities on a Hellinger transformed count table resulting in pair-wise dissimilarity values between all of the individual samples. This is referred to as the dissimilarity object, which was also used in the Adonis analysis, Principal Coordinates Analyes (PCoA), and Procrustes analysis. Statistical tests comparing dissimilarities were performed between the different groups of samples, i.e. within replicates, within the same sample matrix, and between the different sample matrices. A Levene’s test on raw dissimilarities between groups and log transformed dissimilarities revealed that a non-parametric test was appropriate. A Kruskal-Wallis test was used to test whether there was an overall difference between groups and if significant (P<0.05), a post-hoc analysis was performed using a Dunn pair-wise test of groups as part of the FSA package version 0.8.20.

#### Heatmaps

To visualize the taxonomic and resistance gene abundance, heatmaps were generated using the pheatmap package. In the heatmap visualization, microbial genera were standardized to zero mean and unit variance, including the 30 most abundant genera. Sample-based dendograms were generated from the dissimilarity object using complete-linkage clustering. The microorganism-based dendograms were generated using Pearson product-moment correlation coefficients with complete-linkage clustering on the dissimilarity object.

#### Adonis

The ‘adonis2’ function as part of the vegan package was used to perform an analysis of variance. The function takes a dissimilarity object and partitions dissimilarities for the sources of variation. It is a permutational approach and in doing so the significances of the partitions can be assessed. The ‘betadisper’ function was used to assess the differences in group homogeneities and the fitted model were analysed with ‘permutest’. If significant (p<0.05), the model was considered inappropriate for the analysis.

#### Principal coordinate analysis (PCoA)

PCoA plots were generated using the ‘capscale’ function unconstrained as part of the vegan package. The inbuilt stressplot function in vegan was used to extract the ordination distance and corresponding dissimilarities to make stressplots in ggplot2. The coordinates and eigenvalues were extracted from the capscale object to create PCoA plots and screeplots, and visualized with ggplot2. How well the PCoA performed at visualizing the data was evaluated in the stressplots. The screeplots were used to evaluate how much variance was explained on the different axis.

#### Procrustes

Procrustes rotation was performed with the vegan function ‘protest’ to compare the principal coordinates of the taxonomy-based dissimilarities and the antimicrobial resistome-based dissimilarities using classical multidimensional scaling (cmdscale). The descriptive statistics were extracted from the object generated from ‘protest’ and included in the plot made with ggplot2.

#### Canonical correspondence analysis (CCA)

A constrained visualisation of the data was performed with the ‘cca’ function in vegan on the raw count table. CCA is based on the chi square distance and therefore the raw count table was used. The cca plot, screeplots and stressplots were generated from the cca object using ggplot2. The genera in the plot were coloured according to different selected taxonomical groups or antimicrobial resistance (AMR) classes based on occurrence and abundance in the dataset.

#### Differential abundance analysis

Differential abundance analyses were performed using DESeq (31) at sample matrix level (P1, P2, S1 and S2) using the raw count table. Custom DESeq sizeFactors were used taking the total sums for the different samples divided with the mean total sums for all of the samples. Pairwise comparisons of the abundance of the genera were performed between storage conditions using DESeq2. Based on the PCoA plots and raw dissimilarities the difference between 0h vs 64h -80°C, 0h vs 64h 22°C and 0h vs 64h 5°C were further assessed through the use of MA-plots with the genera coloured according to the same criteria as in the CCA (see above). Comparison of the same genera or AMR genes across the different samples was performed by extracting log2-fold changes, and generating boxplots to compare across all samples and scatterplots to compare the two pig feces and sewage samples separately.

#### Mock-Community

Line plots were generated by relativizing to the mean at 0h for spiked and un-spiked samples respectively. All values were log2 scaled and the 95% confidence interval for each measurement is represented in the plots. Stacked bar charts for each storage condition and sample were created for the detected mock community organisms and compared with the microscopy-based cell counts, colony forming units (CFU) and concentrations from the individual DNA isolations obtained for the individual mock community members used for assembling the mock community. To account for the background in the spiked samples a factor (spiked/unspiked) was calculated from the non-mock community genera present in the samples. This factor was unique for sample and storage condition. The factor was used to multiply mock community genera in the unspiked samples obtaining a more precise estimate of the background in the spiked samples and then subtracted from the according genera in the spiked samples. This procedure was performed to account for the compositionality of data and that the mock community recruits reads from the background.

## Data availability

The raw reads are accessible from the International Nucleotide Sequence Database Collaboration (INSDC) at ENA/EBI, DDBJ, NCBI under Project accession PRJEB31650. Additional data are available from Figshare at https://figshare.com/projects/Standard_sample_storage_conditions_impact_on_inferred_microbiome_co mposition_and_antimicrobial_resistance_patterns/76044.

## Acknowledgements

We thank Berit Knudsen (DTU, Denmark), Marie Stengaard Jensen (DTU, Denmark), Carsten Bistrup (DTU, Denmark) and Lyngby-Taarbek Forsyning (Denmark) for assistance with sample collection. We are grateful to Ulrich Nübel (DSMZ) and Sabine Gronow (DSMZ) for providing strain *Propionibacterium freudenrichii* DSM 20271. Oksana Lukjancenko (DTU, Denmark), Patrick Munk (DTU, Denmark), and other members of the GenEpi group (DTU, Denmark) are thanked for their support with the bioinformatics and statistical analysis. The data analysis described in this study was performed using the DeiC National Life Science Supercomputer at DTU (Denmark).

This study has received funding from the European Union’s Horizon 2020 research and innovation programme under grant agreement no. 643476 (COMPARE) and The Novo Nordisk Foundation (NNF16OC0021856: Global Surveillance of Antimicrobial Resistance).

CP and SJP conceived and designed the experiments. CP Performed the experiments. CP and RK analysed the data. CP and SJP performed the literature review and wrote the paper. RK and FA revised the manuscript. All authors read and approved the manuscript.

## Supplemental Figures

**Figure S1:**
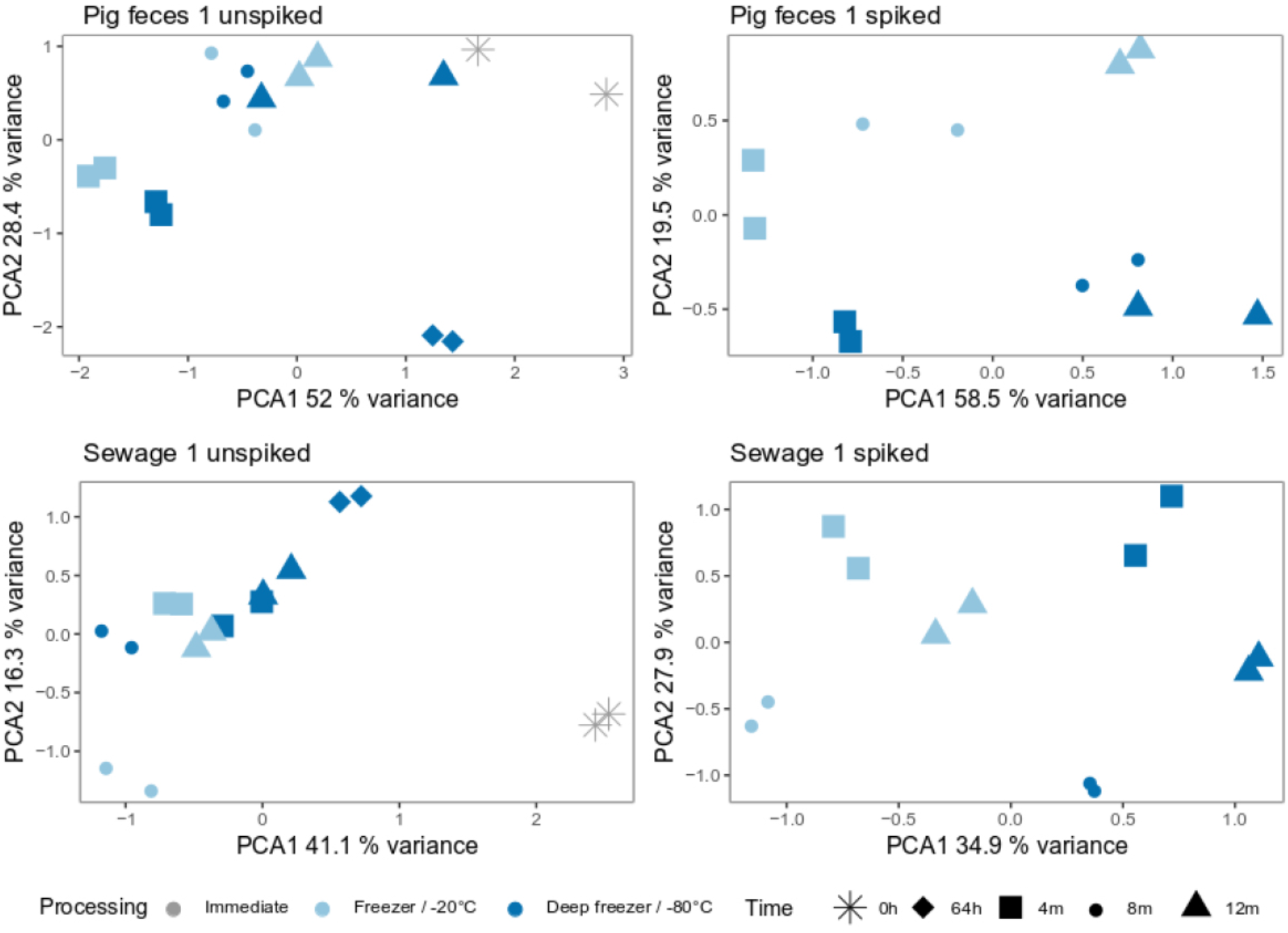
The effect of long-term storage. Principal component analysis (PCA), for pig feces 1 (unspiked/spiked with mock community) and sewage 1 (unspiked/spiked with mock community), to explore the effect of freeze-thaw cycles. PCA were generated from an Euclidean distance matrix obtained from isometric log-ratio transformed data. Variance explained by the first two dimensions are indicated at the axes.

**Figure S2:**
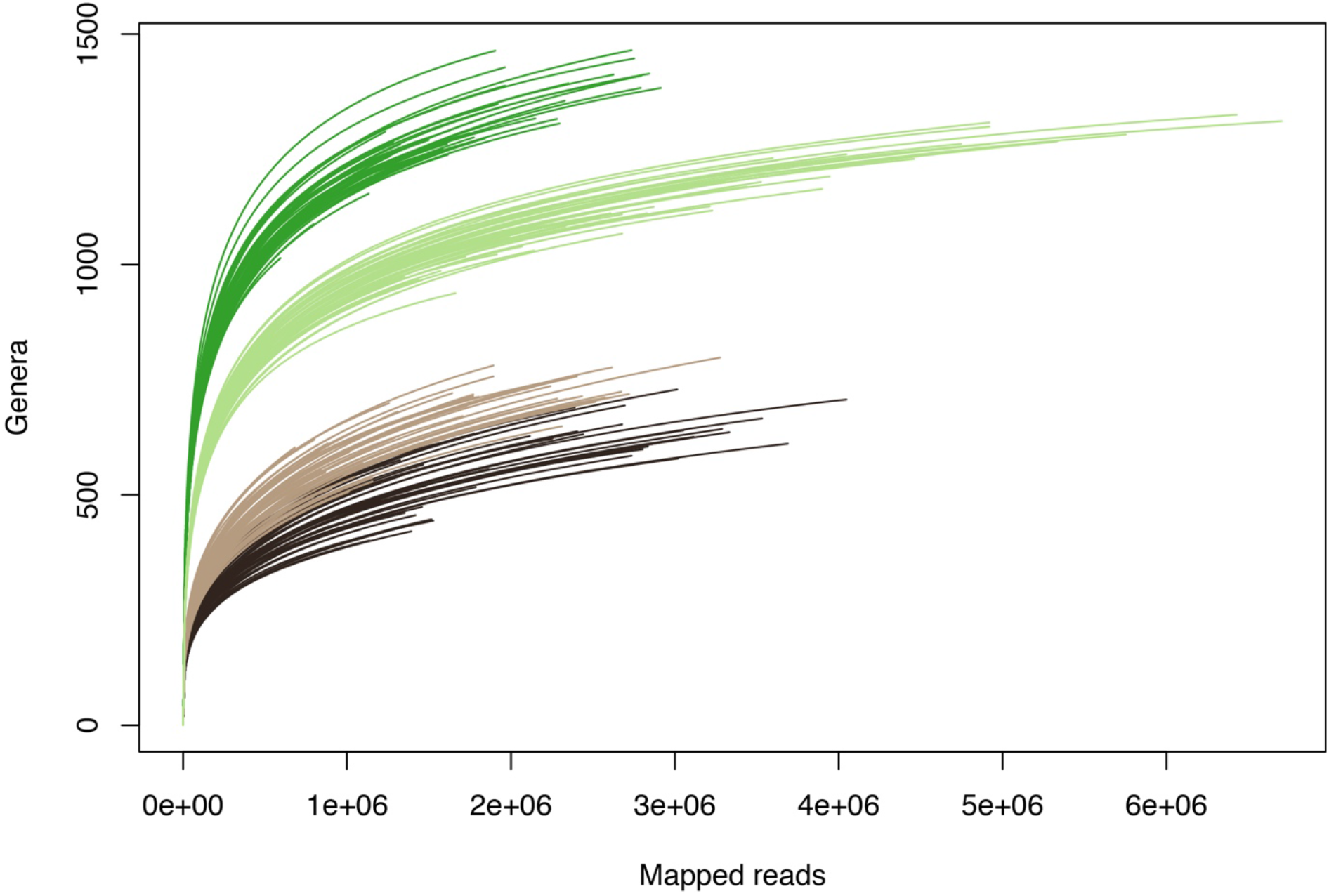
Rarefaction curves for all samples from pig feces P1 (dark brown), pig feces P2 (light brown), sewage S1 (dark green), and sewage S2 (light green).

**Figure S3:**
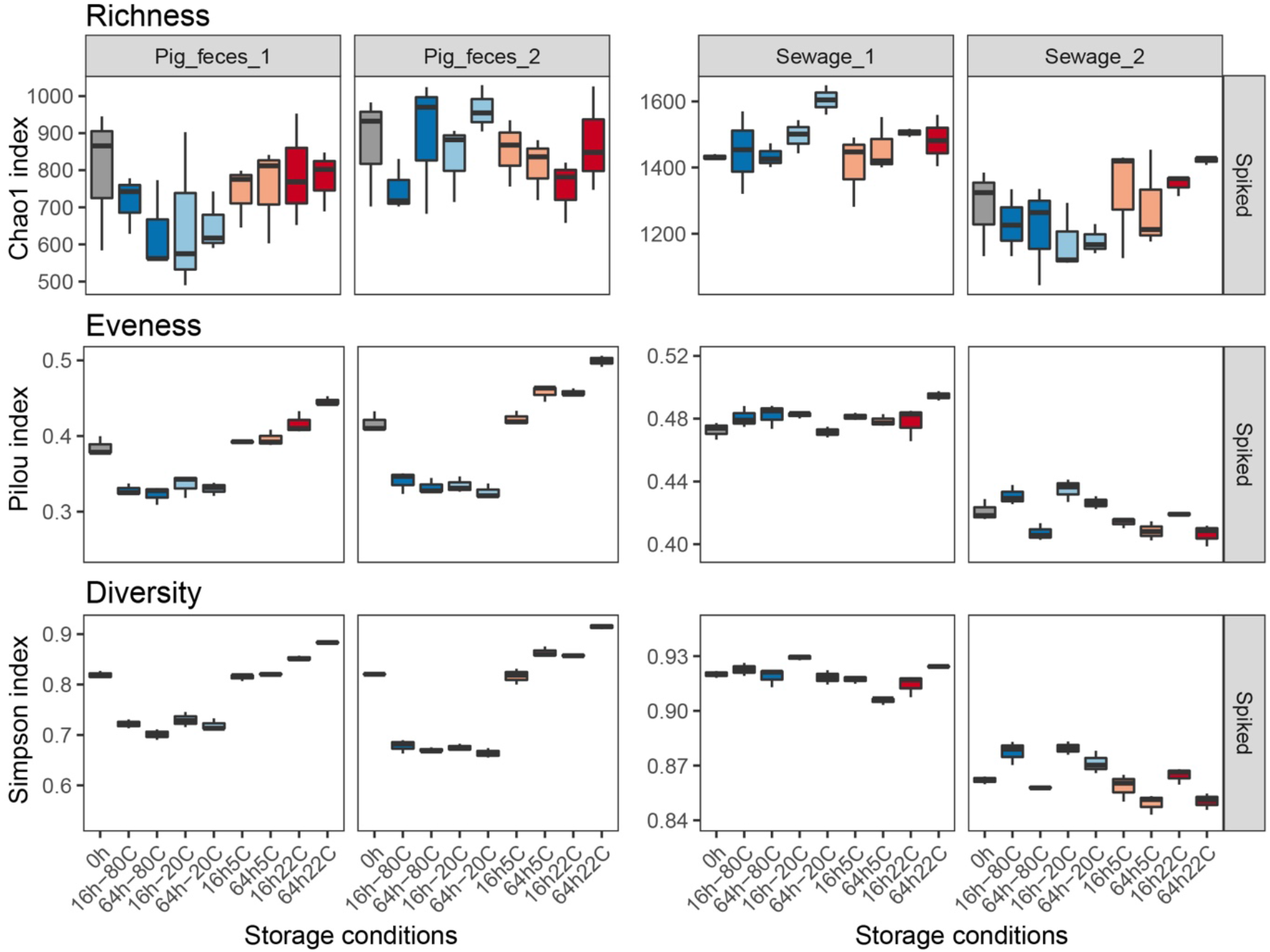
Alpha diversity for samples spiked with a mock community: richness (Chao1), evenness (Pielou’s evenness), and diversity (Simpson). The indices were calculated from the count table aggregated at genus level. For the results of the individual unspiked samples, see Figure S1.

**Figure S4:**
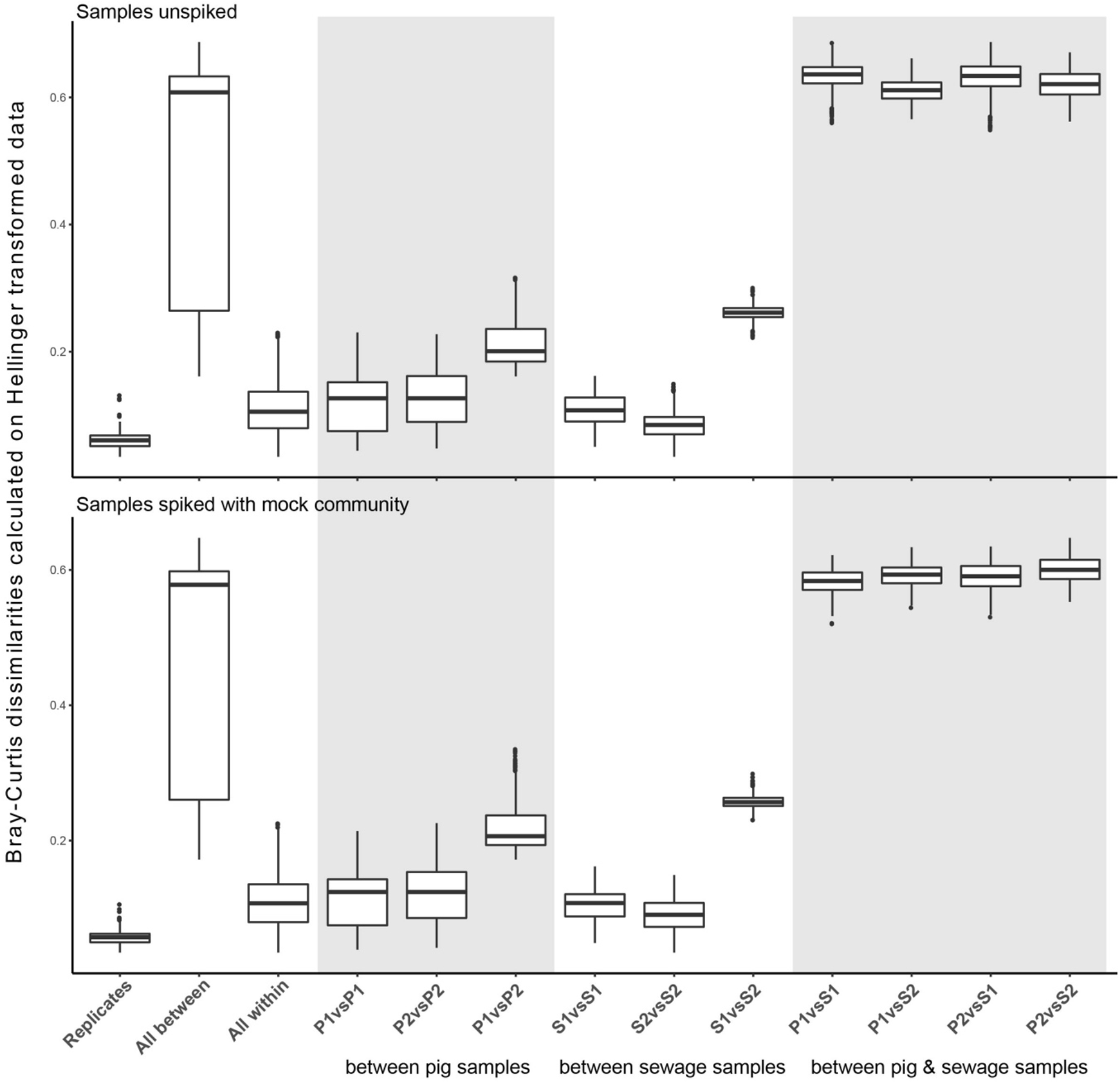
Boxplots of Bray Curtis dissimilarities calculated on Hellinger transformed data. Dissimilarities were grouped according to within sample comparisons and between all possible sample comparison combinations, for unspiked and spiked samples, respectively. Additional boxplots were made for all within and between sample comparisons as well as comparisons between replicates.

**Figure S5:**
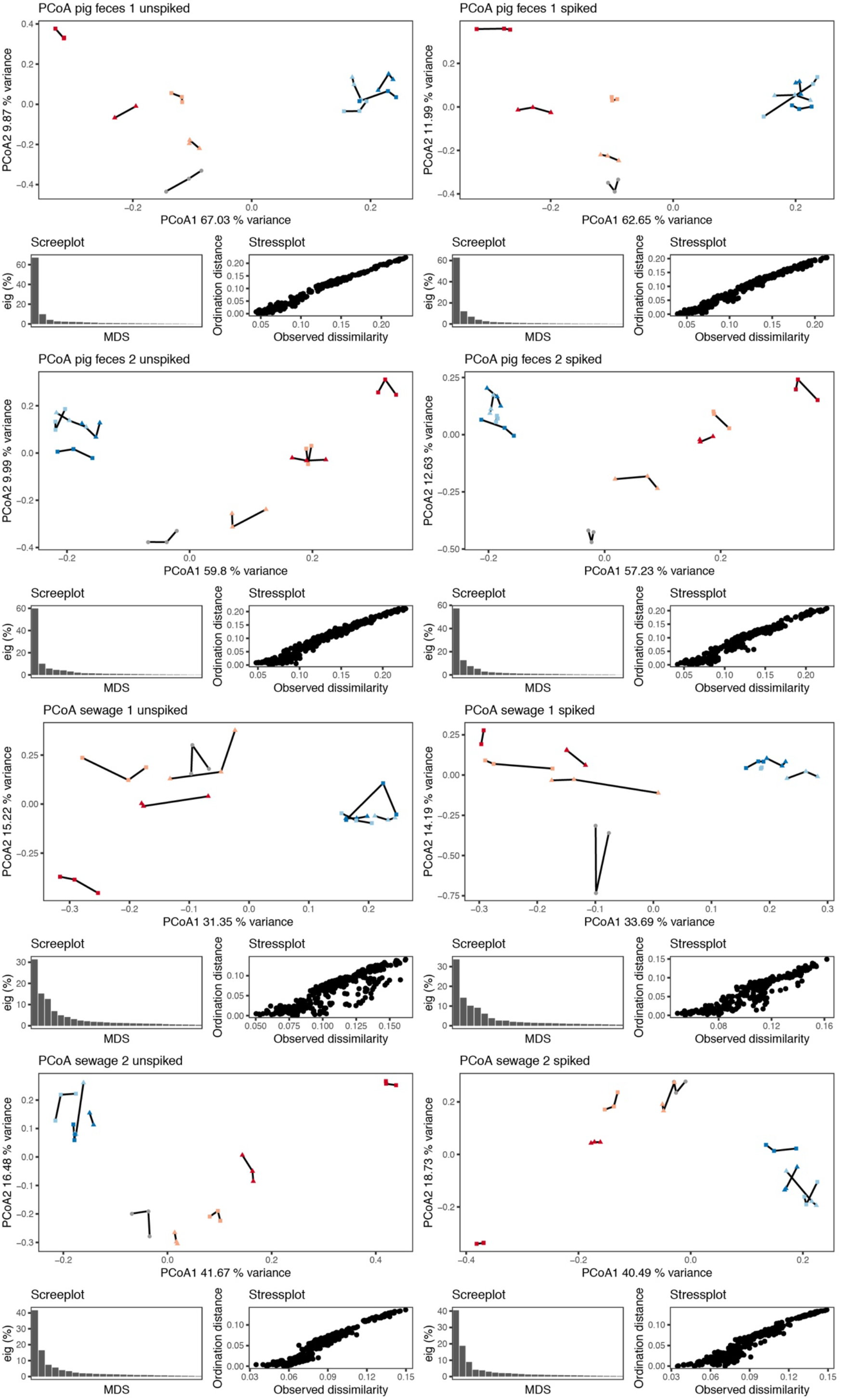
Principal coordinates analysis (PCoA) for each individual sample both unspiked and spiked. Bray-Curtis dissimilarities were calculated on Hellinger transformed data. The vegan function capscale was used to perform PCoA on the dissimilarity matrix. Variance explained by the first two dimensions are included at the axes, respectively. Validation plots (screeplots and stressplots) are included below for each of the corresponding PCoA plot, respectively.

**Figure S6:**
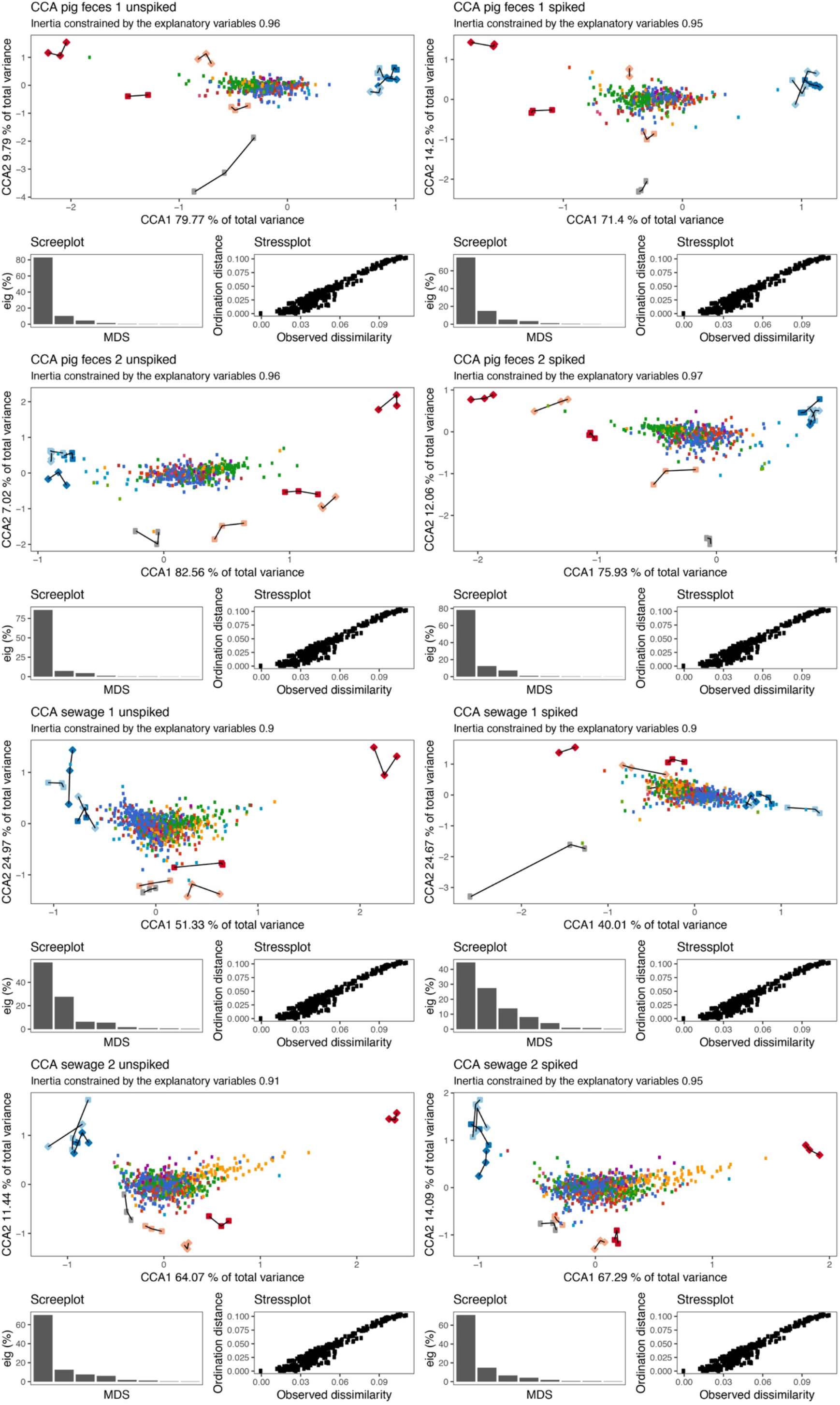
Canonical correspondence analysis (CCA) of each individual samples both unspiked and spiked exploring the taxonomic microbial community composition (coloured points) constrained by the storage conditions (coloured shapes; replicates are connected with lines). The vegan function cca was used to perform CCA on the count matrix. The variance explained by the first two dimensions is included at the axes, respectively. Validation plots (screeplots and stressplots) are included below for each of their corresponding CCA plot, respectively.

**Figure S7:**
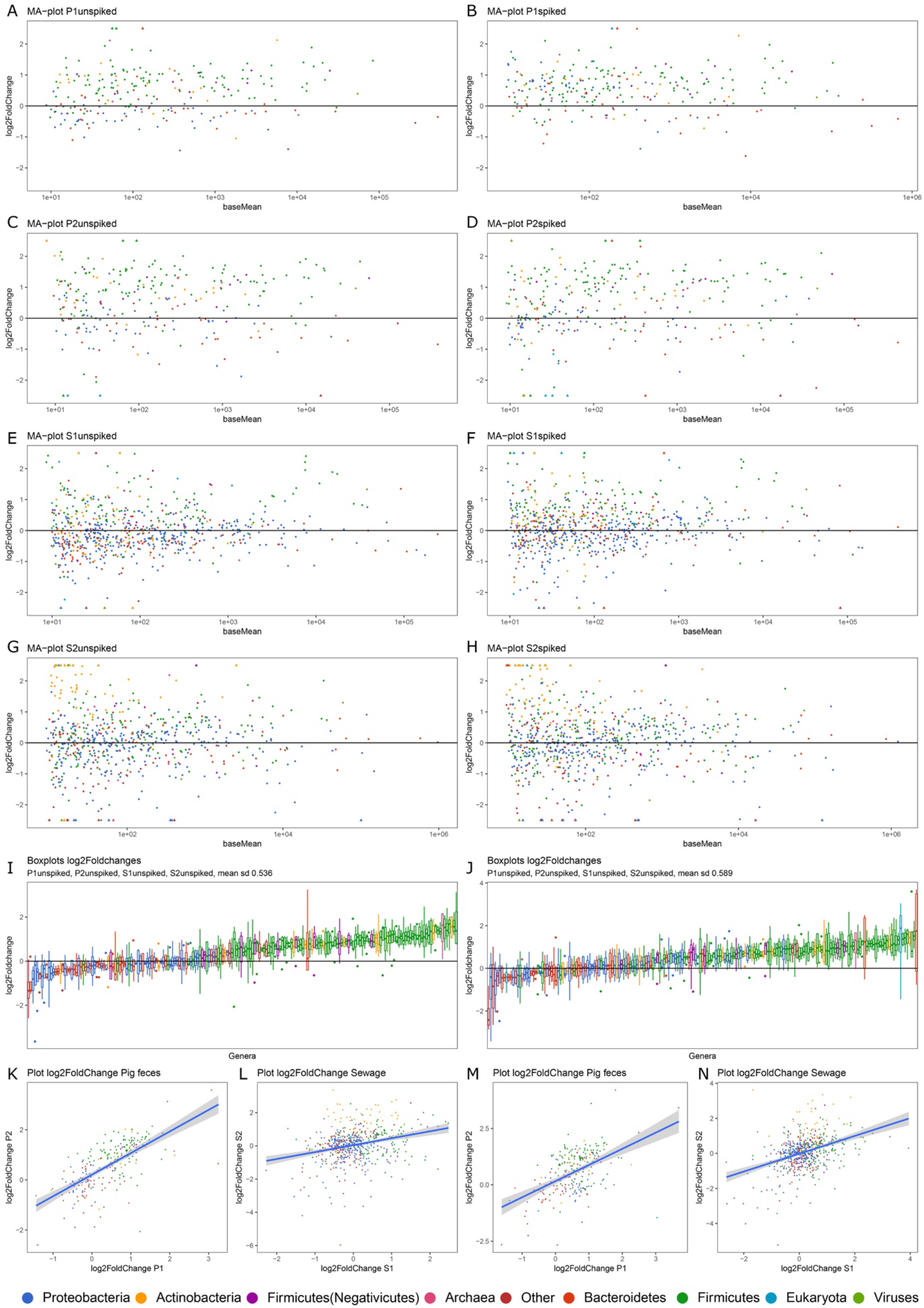
Taxonomic microbial community patterns comparing two storage conditions - immediate DNA extraction & storage at 22°C for 64h. The MA-plots are indicating differentially abundant genera (A-H). Boxplots of log2-fold changes of all shared organisms in all samples sorted from lowest to highest mean (I and J). Scatterplots of log2-fold changes of all shared genera between the two pig fecal samples and two sewage samples (K-N). Plots are generated for both unspiked and spiked sets of samples. Positive log2-fold changes specify a higher relative abundance in samples stored at 22°C for 64h relative to 0 h (A-N). The colour code for the taxonomic groups is indicated at the bottom.

**Figure S8:**
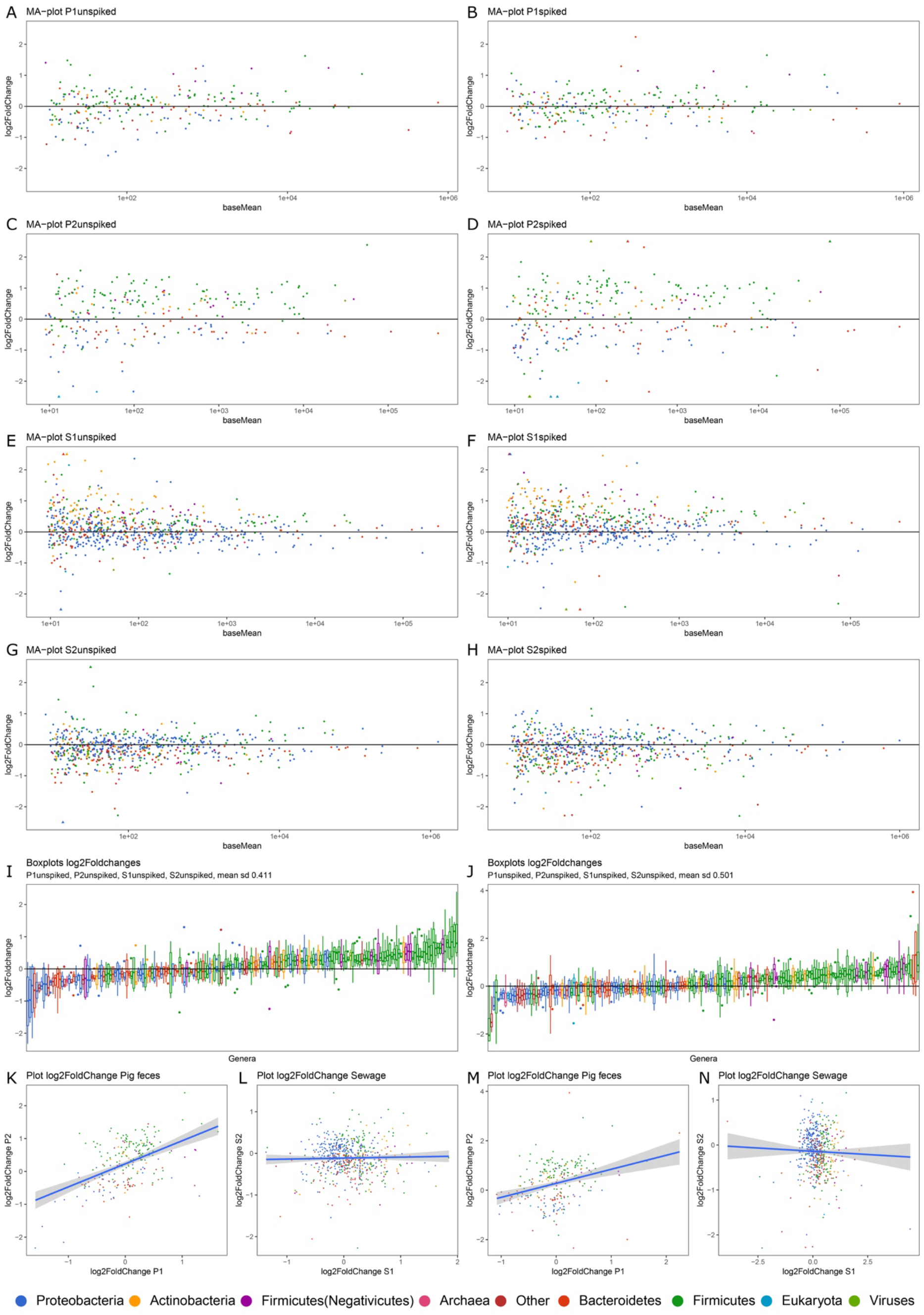
Taxonomic microbial community patterns comparing two storage conditions - immediate DNA extraction & storage at 5°C for 64h. The MA-plots are indicating differentially abundant genera (A-H). Boxplots of log2-fold changes of all shared organisms in all samples sorted from lowest to highest mean (I and J). Scatterplots of log2-fold changes of all shared genera between the two pig fecal samples and two sewage samples (K-N). Plots are generated for both unspiked and spiked sets of samples. Positive log2-fold changes specify a higher relative abundance in samples stored at 5°C for 64h relative to 0 h (A-N). The colour code for the taxonomic groups is indicated at the bottom.

**Figure S9:**
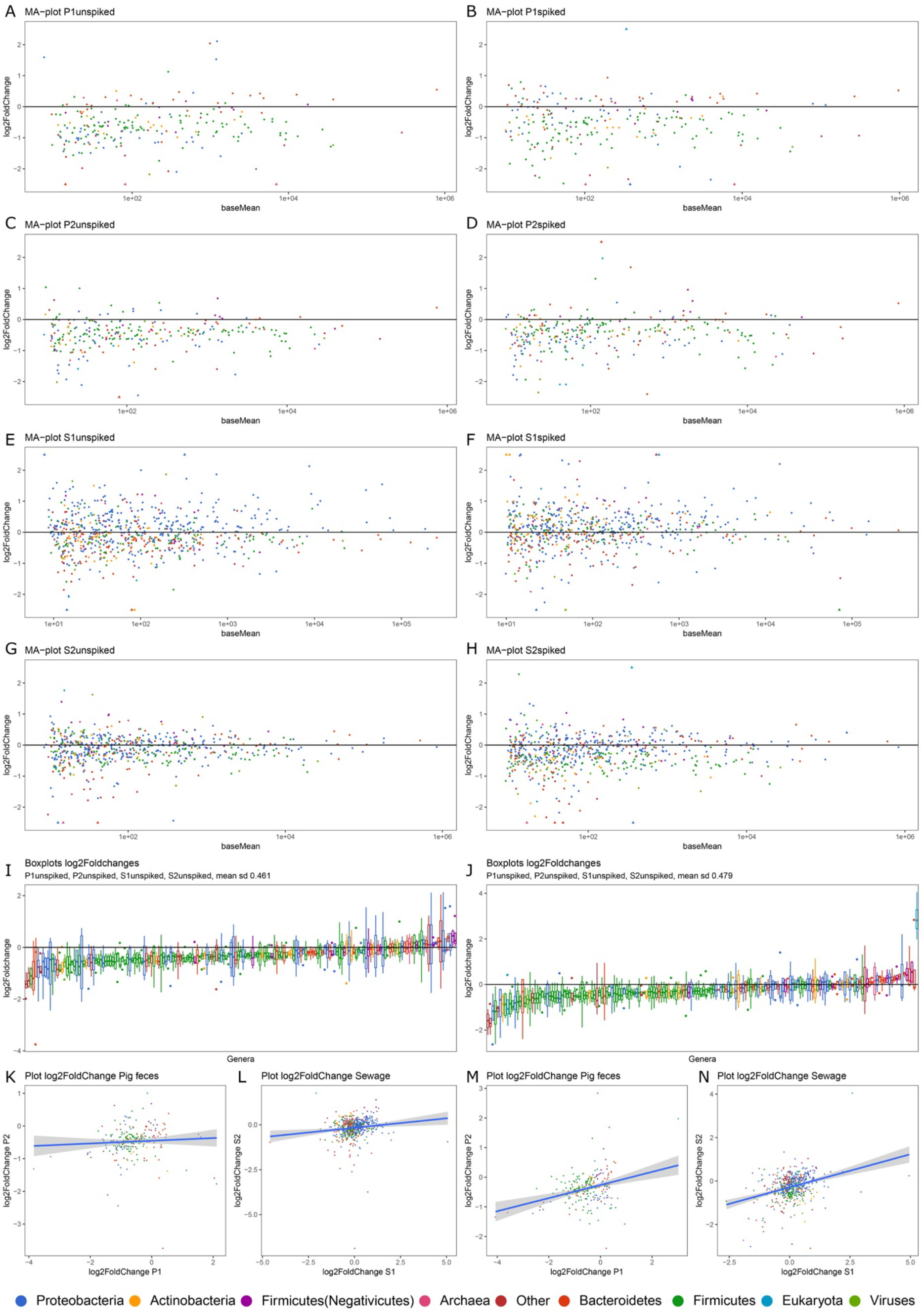
Taxonomic microbial community patterns comparing two storage conditions - immediate DNA extraction & storage at -80°C for 64h. The MA-plots are indicating differentially abundant genera (A-H). Boxplots of log2-fold changes of all shared organisms in all samples sorted from lowest to highest mean (I and J). Scatterplots of log2-fold changes of all shared genera between the two pig fecal samples and two sewage samples (K-N). Plots are generated for both unspiked and spiked sets of samples. Positive log2-fold changes specify a higher relative abundance in samples stored at -80°C for 64h relative to 0 h (A-N). The colour code for the taxonomic groups is indicated at the bottom.

**Figure S10:**
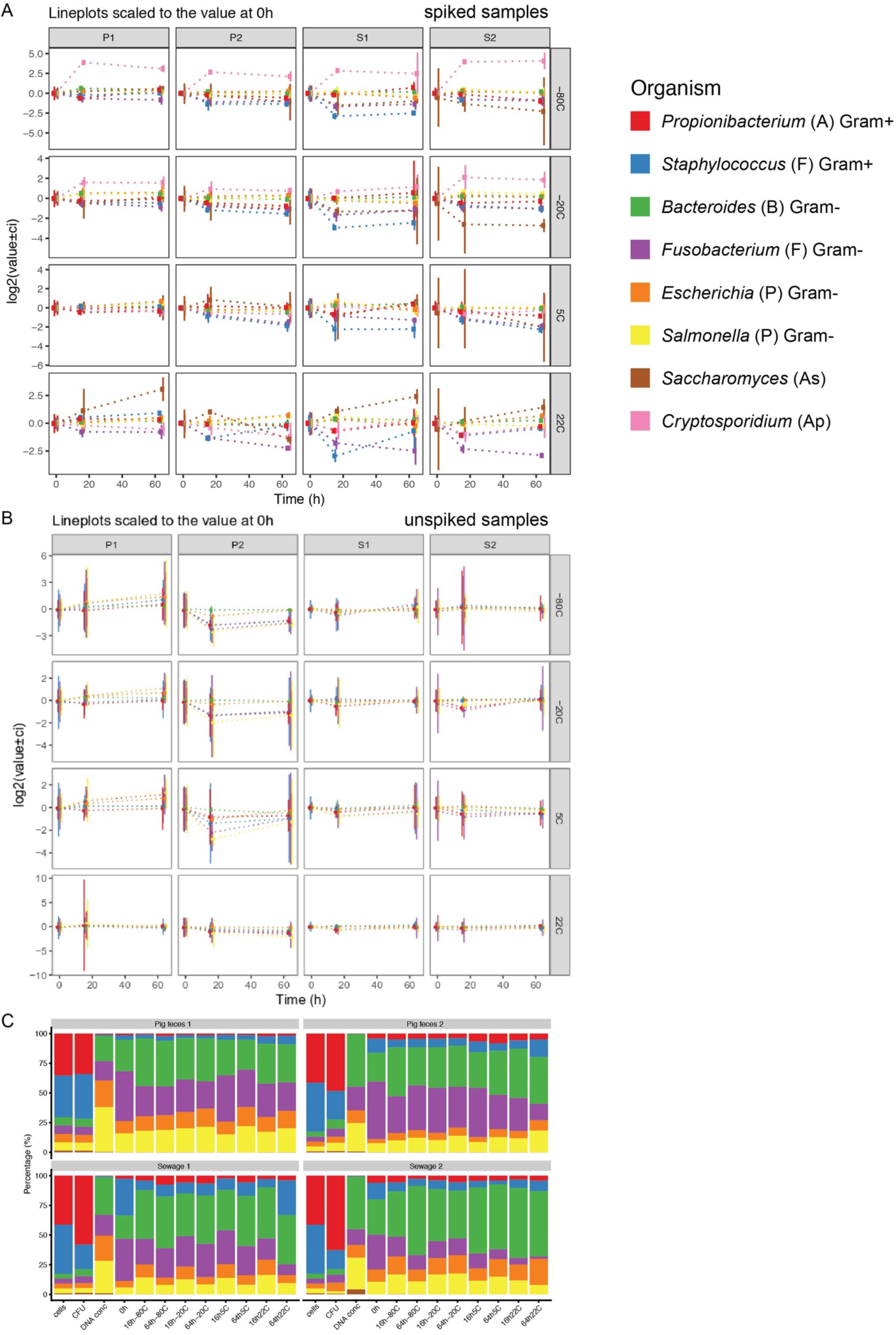
Abundance of the mock community members. (A+B) Lineplots of the mock community organisms in spiked (A) and unspiked (B) samples. The mock community members are represented at genus level namely *Propionibacterium, Staphylococcus, Bacteroides, Fusobacterium, Escherichia, Salmonella, Saccharomyces*, and *Cryptosporidium*. The values were scaled to the average abundance obtained at timepoint 0h. (C) In the first three columns, bacterial cell counts, colony forming units (CFU) and DNA concentration represent the compositions estimated from all cells counted by microscopy in a Petroff counting chamber, viable cells via counting colony-forming units through cultivation, and DNA concentration of the DNA extracts obtained from the individual organisms using the DNA isolation protocol for microbiome samples. The remaining stacked bars represent the microbial community composition at the different storage conditions based on read counts. The read counts represent averages normalized according to genome size and from which the background levels of reads were removed using the information from the unspiked samples.

**Figure S11:**
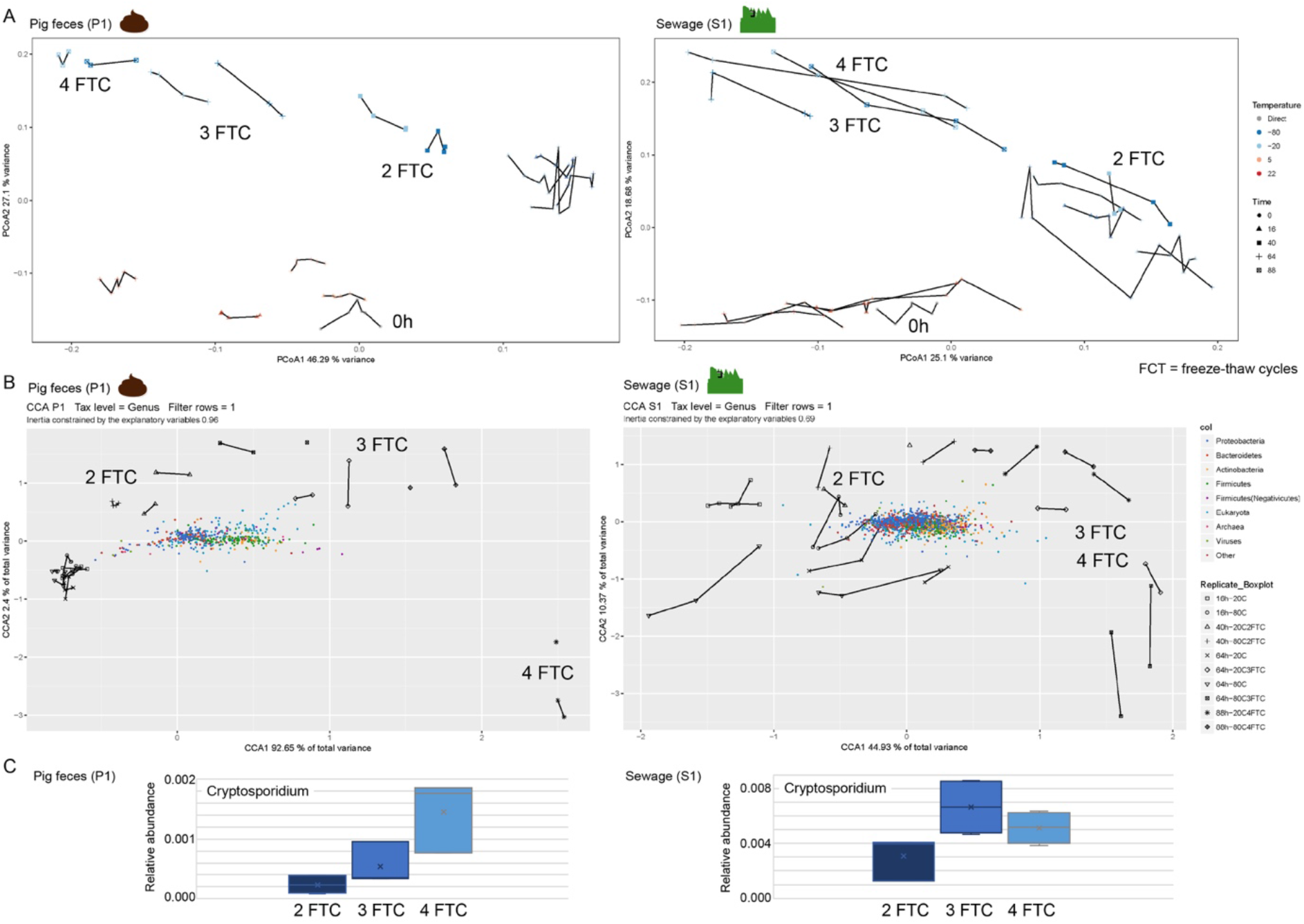
Microbiome patterns upon repeated freeze-thaw cycles. (A) Principal coordinates analysis (PCoA) for pig feces P1 and sewage S1. Bray-Curtis dissimilarities were calculated on Hellinger transformed data. The vegan function capscale was used to perform PCoA on the dissimilarity matrix. Variance explained by the first two dimensions are included at the axes, respectively. (B) Canonical correspondence analysis (CCA) of pig feces P1 and sewage S1 exploring the taxonomic microbial community composition (coloured points) constrained by the storage conditions (coloured shapes; replicates are connected with lines). The vegan function cca was used to perform CCA on the count matrix. The variance explained by the first two dimensions is included at the axes, respectively. (C) The relative abundance of Cryptosporidium upon series of freeze-thaw cycles. The number of freeze-thaw cycles is indicates in the plots.

**Figure S12:**
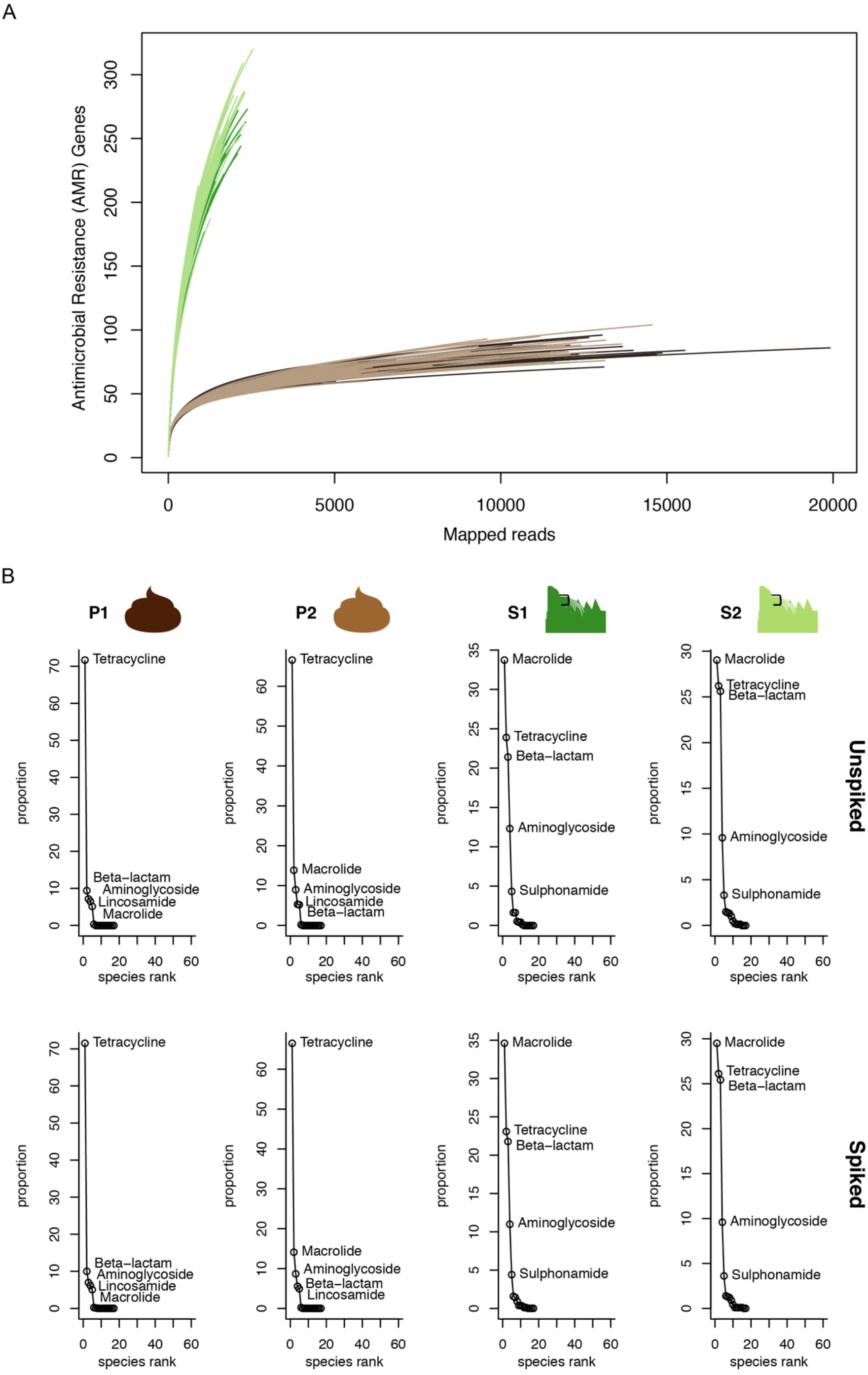
Antimicrobial resistance (AMR) abundance. (A) Rarefaction curves for AMR genes. (B) Rank abundance curves of resistance gene classes showing their abundance distribution in the different samples for both unspiked and spiked pig feces (P1, P2) and sewage (S1, S2).

**Figure S13:**
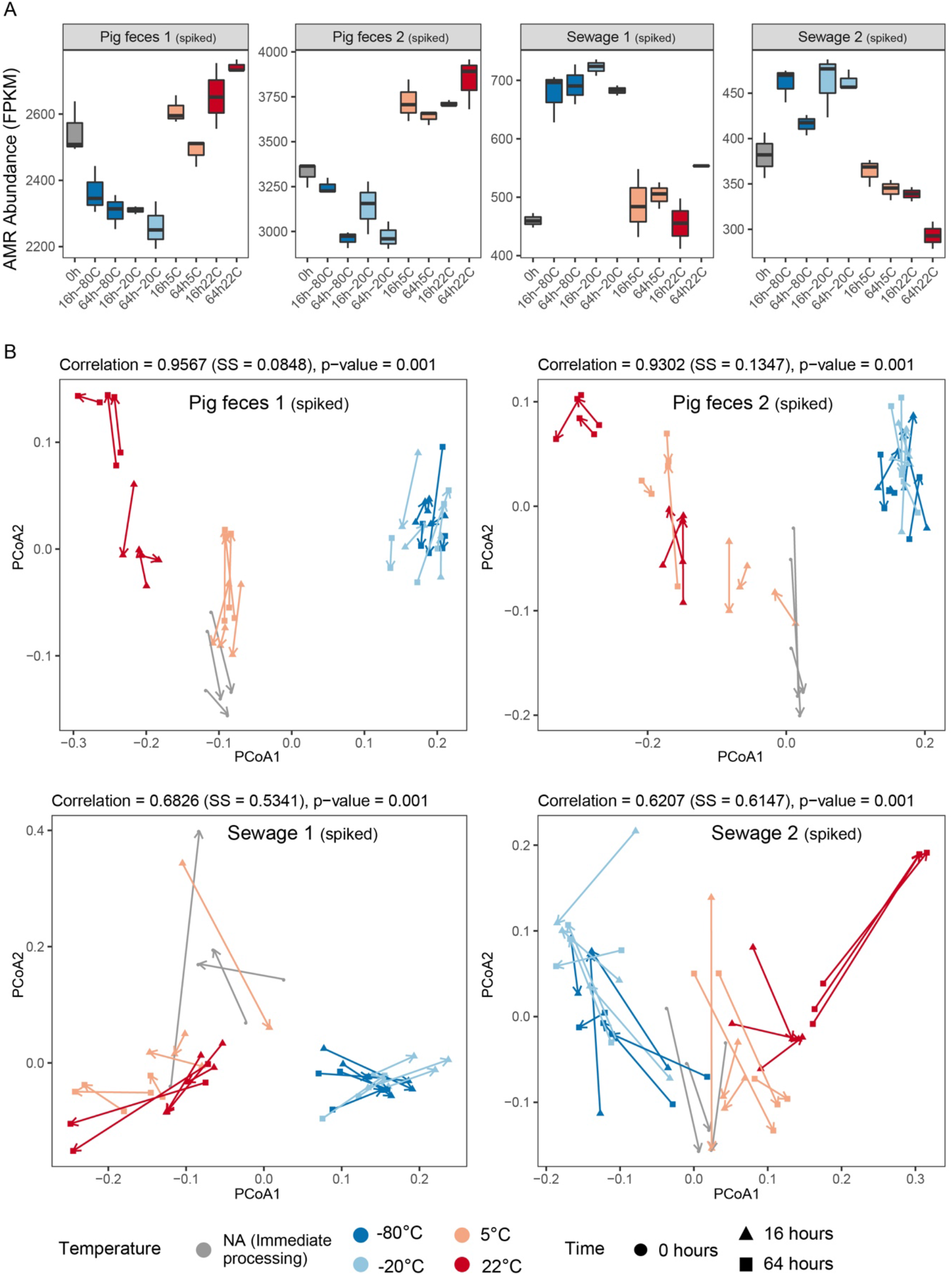
The effect of sample storage conditions on the resistome. A) Boxplots displaying total antimicrobial resistance (AMR) abundance in the spiked samples (P1, P2, S1, S2) measured in FPKM relative to the total number of bacterial reads. B) Procrustes rotation comparing the resistome and taxonomic dissimilarities in the four spiked samples. For the results of the unspiked samples, see Figure 5 in the main text.

**Figure S14:**
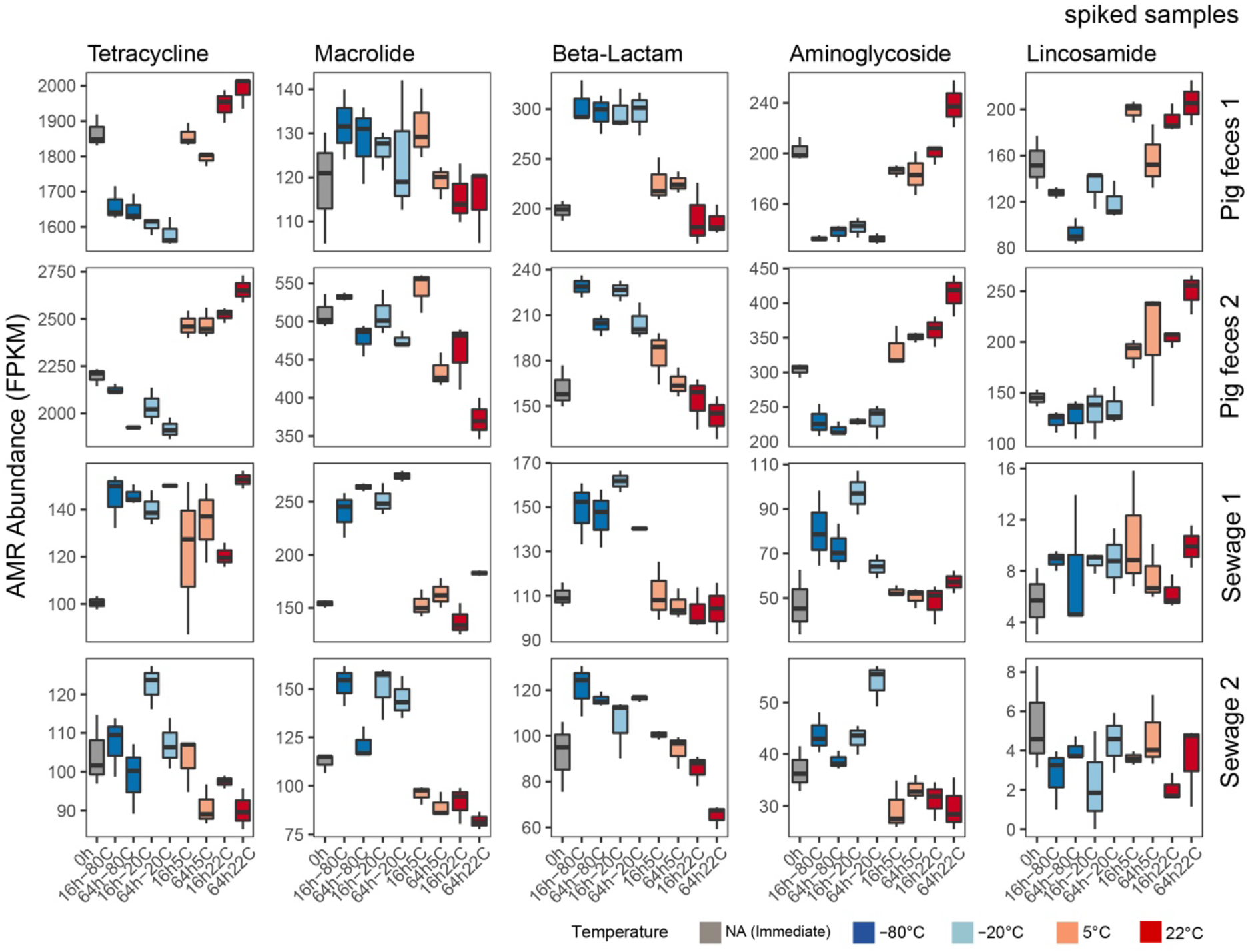
The effect of storage on the abundance of antimicrobial resistance classes. Boxplots displaying total antimicrobial resistance (AMR) abundance for the most abundant antimicrobial resistance classes in the spiked samples (P1, P2, S1, S2). The abundance was measured in FPKM relative to the total number of bacterial reads. For the results of the unspiked samples, see Figure 6 in the main text.

